# Aggregation and immobilisation of membrane proteins interplay with local lipid order in the plasma membrane of T cells

**DOI:** 10.1101/2020.12.22.422352

**Authors:** Iztok Urbančič, Lisa Schiffelers, Edward Jenkins, Weijian Gong, Ana Mafalda Santos, Falk Schneider, Caitlin O’Brien-Ball, Mai Tuyet Vuong, Nicole Ashman, Erdinc Sezgin, Christian Eggeling

**Affiliations:** MRC Weatherall Institute of Molecular Medicine, University of Oxford, Oxford, UK; Jožef Stefan Institute, Ljubljana, Slovenia; Science for Life Laboratory, Karolinska Institutet, Solna, Sweden; Institute of Applied Optics and Biophysics, Friedrich-Schiller-University Jena, Jena, Germany; Leibniz Institute of Photonic Technology e.V., Jena, Germany

## Abstract

The quest for understanding of numerous vital membrane-associated cellular processes, such as signalling, has largely focussed on the spatiotemporal orchestration and reorganisation of the identified key proteins, including their binding and aggregation. Despite strong indications of the involvement of membrane lipid heterogeneities, historically often termed lipid rafts, their roles in many processes remain controversial and mechanisms elusive. Taking activation of T lymphocytes as an example, we here investigate membrane properties around the key proteins – in particular the T cell receptor (TCR), its main kinase Lck, and phosphatase CD45. We determine their partitioning and co-localisation in passive cell-derived model membranes (i.e. giant plasma-membrane vesicles, GPMVs), and explore their mobility and local lipid order in live Jurkat T cells using fluorescence correlation spectroscopy and spectral imaging with polarity-sensitive membrane probes. We find that upon aggregation and partial immobilisation, the TCR changes its preference towards more ordered lipid environments, which can in turn passively recruit Lck. We observe similar aggregation-induced local membrane ordering and recruitment of Lck also by CD45, as well as by a membrane protein of antigen-presenting cells, CD86, which is not supposed to interact with Lck directly. This highlights the involvement of lipid-mediated interactions and suggests that the cellular membrane is poised to modulate the frequency of protein encounters according to their aggregation state and alterations of their mobility, e.g. upon ligand binding.

## 1 Introduction

Cellular plasma membrane hosts numerous vital processes, synchronously executed by a multitude of proteins, many of which are embedded in the fluid lipid bilayer. Once thought of as a homogeneous matrix, it has become clear that this mix of various lipid species and proteins are prone to segregate in more tightly or loosely packed patches, enriched by lipids with saturated or unsaturated alkyl chains and cholesterol, respectively, differing in local lipid order, thickness and molecular mobility (*1, 2*). Employing model membrane systems that exhibit large-scale binary phase separation of lipids, often termed as liquid-ordered (Lo) and -disordered phase (Ld), further investigations revealed that membrane proteins prefer one or the other type of environment based on the structural properties of their transmembrane domains, such as its length, surface area, and lipidation (*3, 4*). Membrane proteins therefore remodel their local lipid environment accordingly, which can supposedly mediate protein-protein interactions, favouring those between proteins in patches of similar lipid composition, order and thickness (*5*). Nevertheless, it has often been notoriously difficult to experimentally dissect this subtle interplay in protein reorganisation observed in trafficking- and signalling-related processes (*6, 7*), for example during activation of T lymphocytes (*8*–*10*).

Activation of T cells, a key element of our adaptive immune response, has been studied extensively to reveal the cellular machinery behind the recognition of foreign antigens (*8*). The very first step in T-cell activation relies on spatiotemporal orchestration of a few key proteins on the plasma membrane: upon binding of the ligand to the T-cell receptor (TCR), immunoreceptor tyrosine-based activation motifs (ITAM) on the TCR-CD3 complex are phosphorylated by a Src family tyrosine kinase (Lck), triggering the signalling cascade downstream. To explain the shift in phosphorylation balance, required for the activation, several models have been proposed. The kinetic segregation model (*11*) advocates that in a resting cell, such activation is prevented by constant dephosphorylation of these ITAMs by the phosphatase CD45, which is sterically excluded from the close contact zone with the antigen-presenting cell to elicit activation. Alternative models involve conformational change of the TCR upon ligand binding (*12, 13*) or its association with cholesterol (*14*), and membrane compartmentalisation (*15, 16*).

While the exact mechanism for the triggering of the TCR remains unclear, mounting evidence has accumulated on the involvement of membrane heterogeneities in T cell activation (*7, 10*). The roles of dynamic nanoscale membrane organisation in this and other processes have remained elusive (*1, 2*), though, primarily due to challenges to experimentally access the relevant spatiotemporal regimes. The first insights were provided by biochemical methods, such as the detergent resistance assay, indicating that upon T-cell activation TCR is relocated from detergent-soluble to detergent-resistant membrane regions (more ordered due to high contents of cholesterol and saturated lipids), which hosts Lck but not CD45 (*17*–*19*). However, these methods are notoriously sensitive to experimental conditions (*2*), casting doubt over reliability and relevance of such conclusions.

To complement these results with gentler experimental methods, several imaging-based approaches have been applied. By inducing ordered membrane patches large enough to be discernible with a conventional confocal microscope, these were found to co-localise with TCR and Lck, but not with CD45 (*20, 21*). Similarly, antibody-induced patches of TCR coincided with higher lipid order, reported by spectral shifts of a polarity-sensitive membrane probe Laurdan (*22*), from which they concluded that TCR in resting cells prefers ordered lipid environment, contradicting the detergent solubility observations. Without such artificial aggregation, conflicting associations of TCR with one or the other membrane environment have been reported in activated live T cells (*23*–*27*) using various advanced optical methods, such as Förster resonance energy transfer (FRET), fluorescence lifetime imaging (FLIM), or super-resolution microscopy. These were likely difficult to reconcile due to different conditions employed (imaging live or fixed cells of different types, different resting/activating states (*28*)), and high TCR density in the non-aggregated state of resting T cells. To this end, model membranes had to be used, where the studies predominantly found monomeric TCR in the disordered membrane areas (*29, 30*).

Though the overview above indicates that TCR may change its preference for its membrane milieu from disordered in the resting state to more ordered in the activated state of the T cell, no mechanism has been provided to compellingly explain this switch in the partitioning behaviour. To further elucidate this lipid-protein interplay, we investigated the lipid environment of TCR and other membrane proteins in passive giant plasma-membrane vesicles (GPMVs) and live Jurkat T cells, using spectral imaging with environment-sensitive membrane dyes. Focusing on the effects of protein aggregation and alterations to their mobility, we find that aggregation and immobilisation of any tested membrane protein, including TCR, can locally induce increased lipid ordering, which can enhance interactions with proteins that prefer such environment, e.g. Lck. Though we cannot firmly conclude whether the observed effects participate in or rather result from the activation of a T cell, our insights nevertheless potentiate a general biophysical mechanism for reorganisation of local membrane order, which could also be applicable to related signalling pathways employing similar machinery, e.g. in activation of B cells (*31, 32*) and mast cells (*33, 34*).

## 2 Results and discussion

### 2.1 Partitioning of proteins in phase-separated GPMVs

To investigate which lipid environment proteins taking part in T-cell signalling (TCR, CD45, Lck, CD4, LAT, CTLA-4, and CD28 as well as its ligand CD86 from antigen-presenting cells) prefer based on their structural properties, such as hydrophobicity, length of the transmembrane domain and lipidation (*4*), we expressed their transmembrane domains (tm, and in the case of Lck its membrane-anchoring SH4 domain), fused to a fluorescent protein, in HEK 293T cells, chosen for their high expression capabilities. Only the multi-span protein complex TCR was expressed in Jurkat T cells to ensure proper membrane presentation. From these cells we prepared giant plasma membrane vesicles (GPMVs) (*35*) containing the fluorescent membrane proteins. By cooling the vesicles to 4 °C and adding deoxycholic acid (DCA), we induced large-scale spatial separation of the vesicle’s membrane into more ordered and disordered lipid regions, termed liquid-ordered (Lo) and liquid-disordered (Ld) phases, respectively (*36*), and verified the protein’s co-localisation with the Ld phase marker Atto647N-DPPE (*37*) (Figure 1A). From the intensity values of two-colour confocal images of the equatorial planes of GPMVs (Figure S1), we further calculated the fraction of molecules partitioning into the Lo phase (Figure 1B). While TCR and tmCD45 showed a clear preference for the disordered phase – to the same degree as the Ld marker (DPPE), Lck (SH4) showed a distinct preference for the ordered membrane environment (Figure 1B).

**Figure 1:**
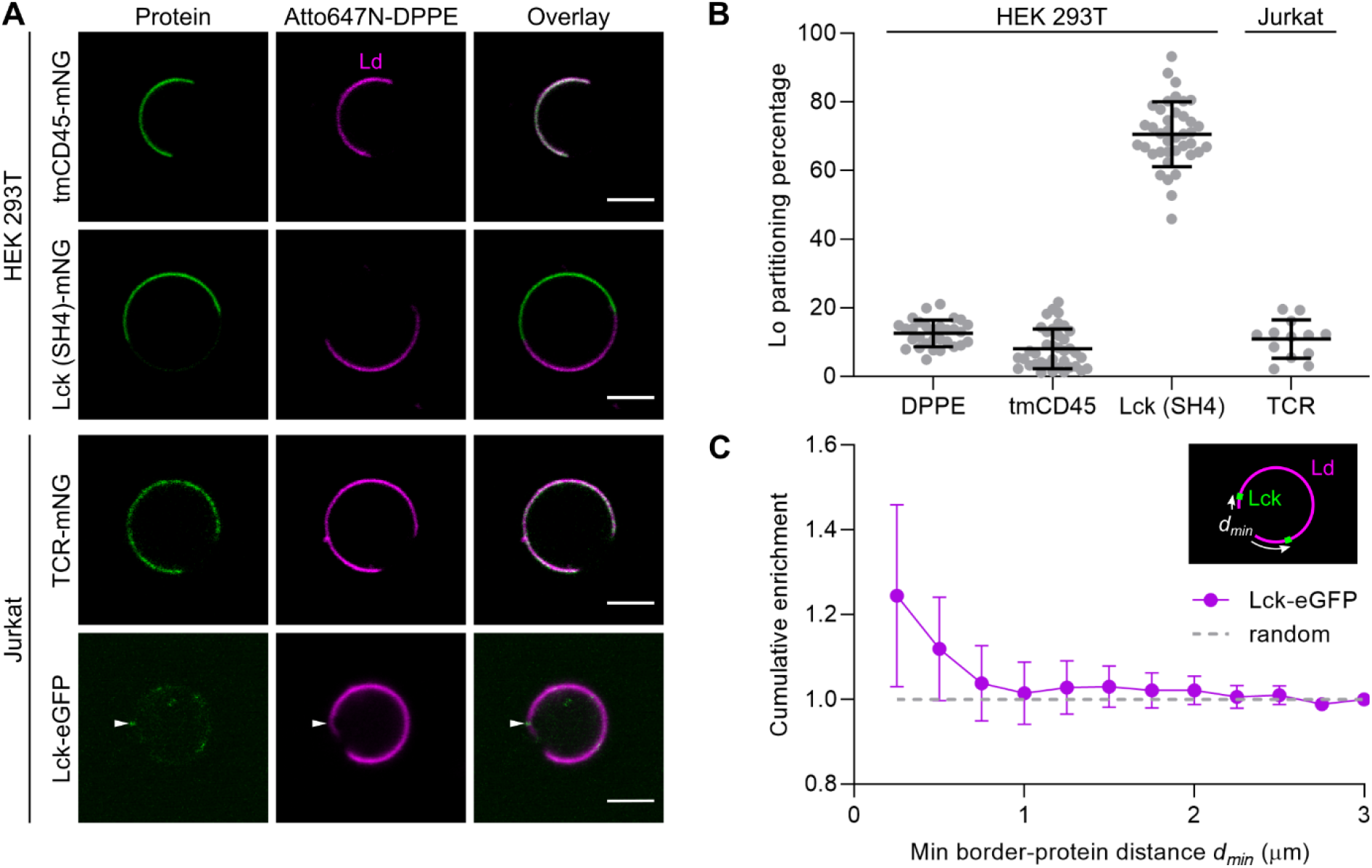
Partitioning of membrane proteins in phase-separated giant plasma-membrane vesicles (GPMVs). (A) Representative images of equatorial planes of GPMVs derived from HEK 293T cells expressing transmembrane (tm) domains of CD45 or membrane-binding Lck domain SH4 fused with mNeonGreen (top left, green), and from Jurkat T cells expressing TCR-mNeonGreen (mNG) or Lck-eGFP (bottom left, green), stained with a liquid-disordered (Ld) phase marker (fluorescent lipid analogue Atto647N-DPPE, magenta, central column, and overlay, right column; scalebar: 5 μm). The arrow in the bottom panel indicates a protein at the border of the lipid phases; see more in Figure S3. (B) Liquid-ordered (Lo) phase partitioning percentages (determined as in Figure S1) for selected proteins and the Ld phase marker (DPPE, control). Every data-point represents a value from a single GPMV, the bars indicate the means and standard deviations. (C) Cumulative enrichment of distances between lipid phase borders and nearest Lck-eGFP (*d*_*min*_, as illustrated in the inset and exemplified in Figure S3C,D) in Jurkat GPMV; values >1 indicate above-random occurrences (see Methods for details).

Our data for Lck (SH4) agree with the published literature, where it has consistently been observed in the ordered lipid environment (*38*–*40*). A similar preference for the ordered lipid environment was displayed also by transmembrane domains of CD4 and LAT (Figure S2A), which are also closely involved in the activation of T cells. Given that partitioning to the Lo phase is most strongly driven by post-translational lipidation of the proteins (*4*), we next verified the roles of these fatty-acid modifications by repeating the experiments expressing Lck (SH4), tmCD4 and tmLAT with mutated cysteine residues, eliminating their palmitoylation sites (Figure S2B–D). For CD4 with a single cysteine residue, the mutation resulted in a complete abolishment of the Lo partitioning. A similar effect was observed for one of the two cysteine sites of LAT in accordance with the literature (*3*), whereas, interestingly, for Lck (SH4) mutation of both cysteine sites induced only a modest reduction in the preference for the Lo phase, likely owing to its additional myristoylation. We obtained consistent results with GPMVs using DTT/PFA, which cleaves off palmitoyl (*3*), but not myristoyl groups (*41*).

Interestingly, the full-length Lck-eGFP construct could not enter the Lo phase in GPMVs of either Jurkat or HEK 293T cells, but was frequently found at the border of the two phases (Figure 1A bottom and Figure S3). Such pronounced enrichment of protein locations at small distances from the phase border (Figure 1C) has previously been observed for some proteins in simulations (*42*) and experimentally (*43*). This behaviour indicates a slight mismatch of the molecule with the disordered lipid phase, and given the artificially dense packing of the ordered phases at the unphysiologically low temperatures used in our model system, one could interpret such border partitioning as a tendency towards the ordered membrane environment, as corroborated by further experiments below.

While the T-cell activating proteins Lck, CD4 and LAT clearly prefer the ordered membrane environment, their key antagonist tmCD45 and the TCR prefer the opposite (Figure 1B), as do transmembrane domains of an inhibitory receptor CTLA-4, a co-stimulatory receptor CD28, and its ligand CD86 (Figure S2A). For TCR, however, conflicting indications have previously been reported: Ld partitioning, as observed here, was reported in model membranes (*29, 30*) and sometimes in resting T cells (*17, 18*), whereas most of the research especially with activated T cells concluded its preference for the highly ordered lipid phase (*17*–*22*).

### 2.2 Partitioning of protein aggregates in phase-separated GPMVs

To resolve this discrepancy, we next isolated GPMVs from Jurkat T cells that had previously been activated by an anti-CD3 antibody (OKT3) in solution, inducing aggregation of the TCR. Interestingly, the induced TCR aggregates were frequently observed stuck to the boundary between the two lipid phases (Figure 2A and Figure S4A), which indicates a change in the preference of the TCR from disordered to a more ordered membrane environment after the activation of the cell and formation of TCR aggregates, in line with previous observations (*17*–*22*).

**Figure 2:**
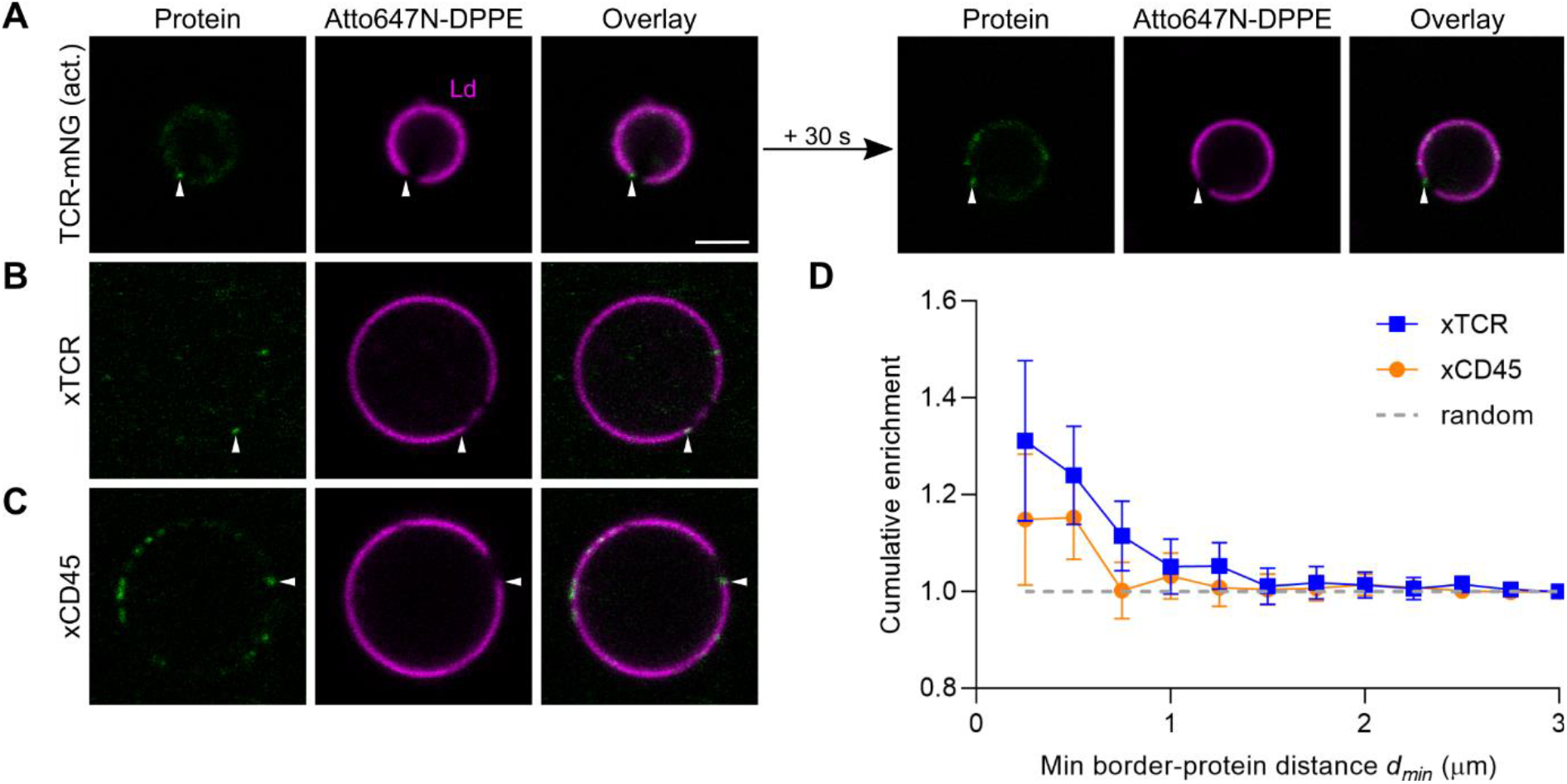
Partitioning of protein aggregates in phase-separated GPMVs. (A–C) Representative images of equatorial planes of GPMVs derived from Jurkat T cells with aggregated proteins (green), stained with the liquid-disordered (Ld) phase marker – fluorescent lipid analogue Atto647N-DPPE (magenta, central column). (A) TCR-mNeonGreen was aggregated in live cells during their activation with OKT3 in solution before the formation of GPMVs. Right: images of the same vesicle 30 seconds later, indicating that protein aggregates stick to the lipid phase border. (B,C) Non-fluorescent proteins were artificially cross-linked (xTCR, xCD45) in GPMVs by incubating with primary (OKT3 against TCR, Gap8.3 against CD45) and fluorescently-labelled secondary antibodies. Arrows point at proteins localised at the border of lipid phases; see more examples in Figure S4. (D) Cumulative enrichment of distances between lipid phase borders and nearest aggregates of TCR (blue) and CD45 (orange) in Jurkat GPMVs; values >1 indicate above-random (grey) occurrences (see Methods for details).

To check whether the change in the lipid phase preference of the TCR upon T-cell activation was caused by an active cellular process or merely as a passive consequence of crosslinking (and thus aggregation), we derived GPMVs from non-activated Jurkat T cells and verified the lipid phase partitioning of TCR cross-linked by primary (OKT3) and secondary antibodies. The induced TCR aggregates frequently partitioned to the lipid phase boundary (Figure 2B,D and Figure S4B), as did those extracted from activated cells (Figure 2A).

Binding of the antibody OKT3 to TCR thus seemingly changed the protein’s preference for the lipid environment, which could result from a conformational change of the protein, or simply from a more general molecular change imposed by aggregation, such as reduced mobility or molecular crowding effects. We therefore examined the partitioning of another Ld-preferring protein, CD45, aggregated with antibodies after isolation of GPMVs – a clearly unnatural condition without a biological function. Interestingly, CD45 aggregates also showed enriched localisation at the border of lipid phases (Figure 2C,D and Figure S4C), highlighting that aggregation of different Ld-partitioning proteins can drive their preference towards more ordered lipid environment in a completely passive way.

We also verified that the effect was not due to a specific property or component of the membranes from T cells: we expressed TCR in Chinese hamster ovary (CHO) cells and therefrom derived GPMVs, in which antibody-aggregated TCR showed similarly frequent border partitioning (Figure S4D).

### 2.3 Membrane order around protein aggregates in GPMVs

In the experiments above, we induced large-scale phase separation in GPMV membranes, hosting almost native complexity of lipid and protein composition (*35*), by cooling the sample to 4 °C and addition of the chemical DCA. Such membrane system is commonly used as an experimentally convenient model believed to mimic physical and chemical properties of small-scale membrane heterogeneities in GPMVs (*44*) and live cells under physiological conditions (*31, 37*). However, the packing densities and diffusion rates were clearly affected by low temperature (*45*) and addition of chemicals (*46*), which could affect the preference of protein aggregates for a specific membrane state.

To verify if any local ordering could be detected at the sites of protein aggregates in seemingly homogeneous (i.e., on the spatiotemporal scales of confocal microscopy) GPMV membranes at room temperature and without DCA, we stained the vesicles with a polarity-sensitive membrane dye NR12S (*47, 48*), which shifts its emission spectrum towards longer wavelengths in more disordered membrane environment. Spectrally resolved imaging (Figure S5A) allowed subsequent unmixing of the overlapping signal contributions from the protein and NR12S as well as determination of its spectral peak position by spectral fitting (Figure S5B) in every pixel of the image (Figure 3A) (*49*). At the sites of both TCR and CD45 aggregates, the fluorescence of NR12S was on average slightly shifted to shorter wavelengths (Figure 3B, Figure S5C), indicating that protein aggregates can locally induce a more ordered lipid environment. A similar effect was detected also for TCR aggregates in GPMVs derived from CHO cells (Figure 3B), again excluding the possibility of the phenomenon being specific to T cells.

**Figure 3:**
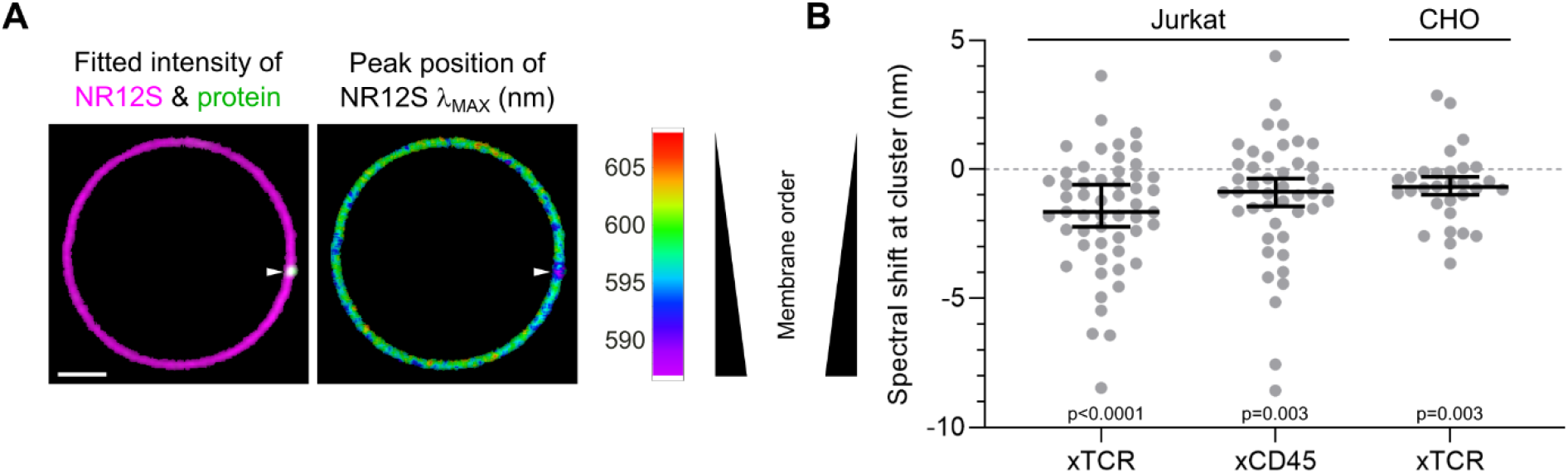
Membrane order in GPMVs with proteins aggregated by antibodies. (A) Spatial maps of the fitted intensities (left) of the protein-aggregating antibody (green; a protein cluster indicated by the white arrow) and NR12S (magenta), and its spectral peak position (λ_MAX_, right; colour scale as depicted: lower values indicate a more ordered membrane environment; scale bar: 2 μm). (B) Spectral shift of NR12S at each identified protein cluster compared to the rest of the membrane, for cross-linked TCR and CD45 in GPMVs from Jurkat T cells, and for TCR aggregates in GPMVs from CHO cells. Two-tailed p-values were calculated by the Wilcoxon sign-rank non-parametric test, against the hypothetical median value of 0. Bars indicate medians and their 95% confidence intervals. For details, see Methods and Figure S5.

The observed 1–2 nm average spectral shift is small, but consistent and thus statistically significant (Figure 3B). One should also bear in mind that the images were recorded with confocal resolution, which likely exceeds the size of the protein aggregates and thereby of the induced regions of increased lipid order, meaning that the real local shifts on smaller scales would be proportionally larger. As the same dye shows a ten-fold larger (i.e., 15 nm) difference between the spectral emission peaks in both phases of large-scale phase-separated GPMVs from Jurkat T cells (Figure S6), we can estimate the average size of the induced more ordered regions to a tenth of the axial confocal cross-section (i.e., 100–150 nm), which is not unreasonable for an aggregate of multiple cross-linked proteins.

### 2.4 Membrane order around immobilised proteins in live T cells

Above, we have used GPMVs as model membranes to investigate the preference for, and influence on, the local membrane environment of membrane proteins. Though GPMVs pertain most of the compositional complexity of the plasma membrane (*35*) and are thus believed to be the most relevant passive mimicking system, certain key structural properties are nevertheless eliminated during their formation, such as inter-leaflet asymmetry (*36*) and interactions with the cytoskeleton (*50*). To verify if our observations translate also to live cells, we next investigated the local lipid order at the sites of the proteins of interest in Jurkat T cells, activated on OKT3-coated surface. Spectral imaging with polarity-sensitive dyes and spectral fitting were again employed to concurrently devise maps of protein density (Figure 4A, central column, green) and membrane polarity, reporting on the local lipid order (Figure 4A, right column). Through the detected spectral shifts of NR12S at the sites of high protein density with respect to the rest of the membrane, variations in the lipid environment of proteins were examined. By filtering of pixels according to the protein intensity relative to the local NR12S intensity, we reduced the influence of bright spots enriched with both protein and lipid signal (e.g. vesicles), allowing us to increase the sensitivity to subtle spectral differences stemming from sub-resolution features (please see Methods for details).

**Figure 4:**
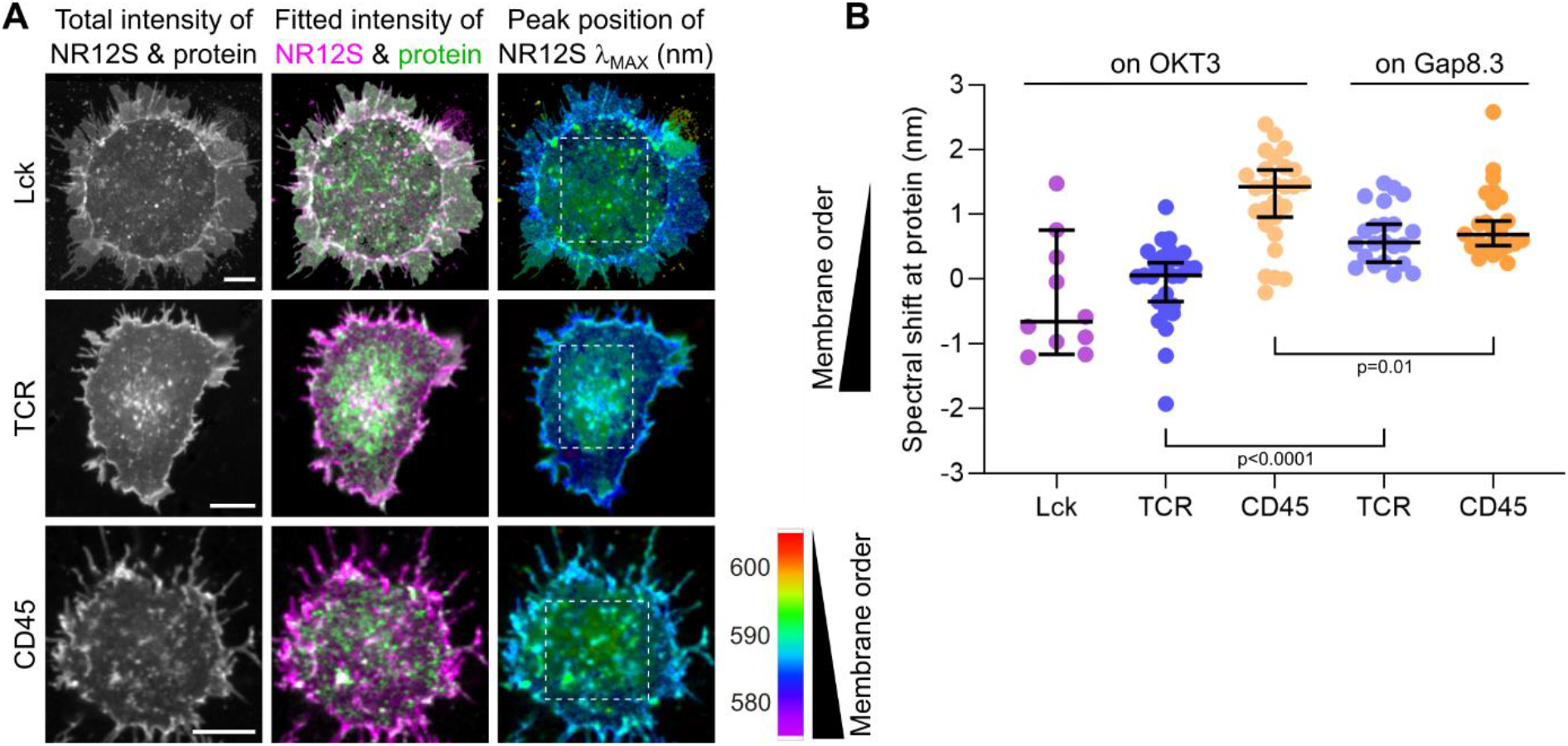
Membrane order in Jurkat T cells on antibody-coated glass surface, probed via spectral imaging with a polarity-sensitive membrane probe NR12S. (A) Representative images of live Jurkat T cells activated on OKT3-coated surfaces: overall intensity (left column), spectrally decomposed intensities of the protein (central column, green; Lck-eGFP (top row), TCR (middle) and CD45 (bottom) labelled with a fluorescent antigen-binding fragment) and NR12S (magenta), and its peak position (λ_MAX_; colour scale as depicted: lower values indicate more ordered membrane). The areas within the white dashed rectangles were used to calculate the (B) relative spectral shifts of the membrane regions with high protein signal, compared to the rest of the membrane for each cell, for Jurkat T cells activated on surface coated with OKT3 (anti-CD3), or Gap8.3 (anti-CD45). Every data-point represents a single cell, the bars indicate medians and their 95% confidence intervals. The p-values were obtained using the two-tailed Mann-Whitney non-parametric test.

In line with the measurements of non-aggregated proteins in GPMVs (Figure 1), Lck and CD45 clearly coincided with more ordered and disordered membrane environments, respectively (Figure 4B). TCR, on the other hand, now showed a tendency towards more ordered membrane regions than CD45 (Figure 4B), corroborating the change of its preference seen upon cell activation or aggregation, observed in the model systems above.

In GPMVs, we observed very similar membrane remodelling by aggregates of both TCR and CD45, potentiating a general passive underpinning molecular mechanism, exerted by protein aggregation. To further verify whether similar membrane ordering effects could be exhibited in live T cells as well, we repeated the last experiment by dropping the cells on a Gap8.3-coated surface to aggregate and immobilise CD45 proteins. In this case, the local membrane order around CD45 increased compared to the previous experiment with an OKT3-coated surface (Figure 4B), whereas the ordering at non-engaging TCR was now slightly released. These results imply that in live cells, aggregation and immobilisation of a membrane protein can locally induce a more ordered membrane environment.

In the experiment above, aggregation and full immobilisation of the protein engaging with surface-bound antibodies was employed, which is certainly a simplistic setting and may have exaggerated the observed effects. In a physiological case, the interaction of TCR and co-receptors with a peptide-loaded major histocompatibility complex (pMHC) on an antigen-presenting cell with a fluid plasma membrane would impose a significant slow-down of the protein aggregate (*51*), but not to a complete stall.

Coming a step closer to a realistic scenario while keeping well controlled conditions and planar geometry suitable for microscopy of the interface, we substituted the antibody-coated surface with a supported lipid bilayer (SLB) – a fluid artificial membrane deposited on a coverslip. The SLBs were decorated with diffusing molecules of CD58 (*52*), which binds to the adhesion protein CD2 on the surface of T cells and thereby reduces their crawling while imaging, but did not promote their spreading (Figure 5A, top row, “adh”). To prompt the engagement of TCR with the SLB, we either incorporated pMHC (Figure 5A, bottom), or labelled TCR with an anti-CD3 antigen-binding fragment (fab) with a large extracellular protein domain (UCHT-1 fused to CD45R0) possessing a His-tag group (Figure 5A, middle row, “TCR-His”), which binds nickel-nitrilotriacetic acid (Ni-NTA)-functionalised lipids in the SLB. None of these conditions would explicitly promote cross-linking of TCR.

**Figure 5:**
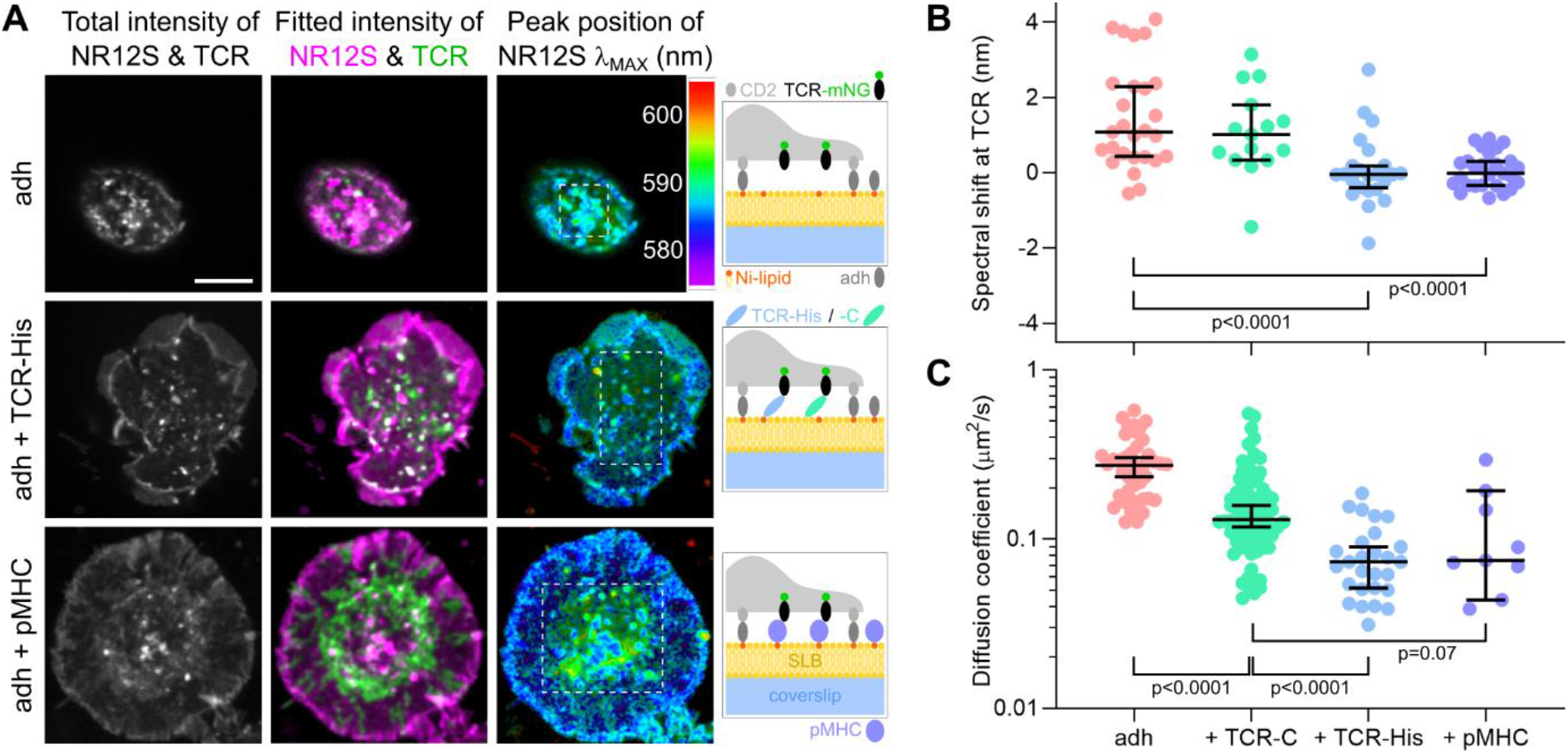
Membrane order and diffusion of TCR in Jurkat T cells on supported lipid bilayers (SLB). (A) Sample images of live Jurkat T cells expressing TCR-mNG: overall intensity (left column; scalebar: 5 μm), spectrally decomposed intensities of the TCR (central column, green) and polarity-sensitive membrane dye NR12S (magenta), and its peak position (λ_MAX_; colour scale depicted, lower in more ordered membrane). Cartoons depict experimental conditions: all SLBs were decorated with the adhesion protein CD58 (adh) to attach the cells for imaging. TCR was made to interact with the SLB via the pMHC complex on the SLB (bottom row), or via an anti-CD3 fab with a His tag (TCR-His) that binds nickelated (Ni-) lipids in the SLB (middle row). The areas within the white dashed rectangles were used to calculate the (B) relative spectral shifts of the membrane regions with high protein signal, compared to the rest of the membrane of each cell. (C) Diffusion coefficient of the TCR-mNeonGreen in its high-density regions in Jurkat cells on SLBs under different conditions (TCR-C: TCR labelled with a C-tag fab, which does not bind to lipids in the SLB), obtained with FCS. Every data-point represents a single FCS recording. In B and C, the bars indicate medians and their 95% confidence intervals. The p-values were obtained using the two-tailed Mann-Whitney non-parametric test.

Compared to the case with no specific TCR engagement with the SLB (“adh”), TCR binding to SLB via both pMHC and TCR-His induced similarly pronounced spreading of the cells (Figure 5A), local ordering of the membrane around TCR (Figure 5B), and slow-down of the diffusion of TCR proteins (Figure 5C), measured by fluorescence correlation spectroscopy (FCS) in regions of the cell with the highest TCR density (e.g. aggregates). A similar fab with the His-tag substituted by a C-tag (“TCR-C”), unable to bind the opposing SLB, slowed down TCR considerably less (Figure 5C) and induced no detectable local membrane ordering (Figure 5B). This shows that an approximately 3–4-fold reduction in diffusivity corresponded to a comparable change in local membrane order as for immobilised or cross-linked proteins (Figure 4B and Figure 3B, respectively).

Noteworthy, both conditions with slow down-induced membrane ordering (i.e. SLB with pMHC and TCR labelled with the His-tag, but not the C-tag fab) also elicited robust T-cell triggering, as probed by cytosolic Calcium release (Chen et al., submitted). Though the increased membrane ordering around TCR is linked with activation of the cells, we cannot firmly determine the direction of causality, as our measurements were performed several minutes after landing of the cells and thus after activation. Nevertheless, our findings with live-cell microspectroscopy importantly corroborate very similar observations with super-resolution microscopy with fixed B cells (*31, 32*), implying common underlying molecular mechanisms. To further evaluate the possibility of involvement of lipid reordering in the activation process, we reverted to a passive membrane model system, as discussed next.

### 2.5 Co-localisation of Lck with protein aggregates in GPMVs

T-cell activation relies on TCR being phosphorylated by Lck. Our results, described above, indicate that in the resting state with both proteins freely diffusing, the frequency of their random encounters could be kept low also by their intrinsic preference for different type of local membrane environment. Upon engagement of TCR (and its co-receptor CD4) with pMHC, leading to the slow-down of the protein aggregate, however, the modified local membrane order resembled more the preferred environment of the Lck, potentially fostering their interaction required for the triggering to occur.

The polarity-sensitive dye used so far, though, can only report on the similarity of the local lipid order of the membrane patches created by protein immobilisation and those hosting Lck, but cannot reveal their identity or interactions. To finally verify whether these lipid heterogeneities can mediate the attraction between the proteins, we investigated co-localisation of Lck with antibody-induced aggregates of TCR in GPMVs – a passive model membrane system devoid of cellular processes. Despite weak signals, owing to low incorporation of Lck into the GPMVs and down-regulation of TCR in the cell line over-expressing fluorescent Lck, we observed many occurrences of its coincidence with TCR aggregates (Figure 6A). This was corroborated also by a quantitative analysis of the extracted two-colour intensity profiles along the membrane (Figure 6B; the Overlap intensity ratio Z score measures the increase in signal co-localisation compared to randomly scrambled intensity profiles, values above 0 indicate an above-random degree of co-localisation; see Methods for details). We verified that the aggregates did not attract all the membrane components in an unspecific manner, as the membrane dye DPPE did not show any enrichment at the sites of protein aggregates (Figure 6B and Figure S8).

**Figure 6:**
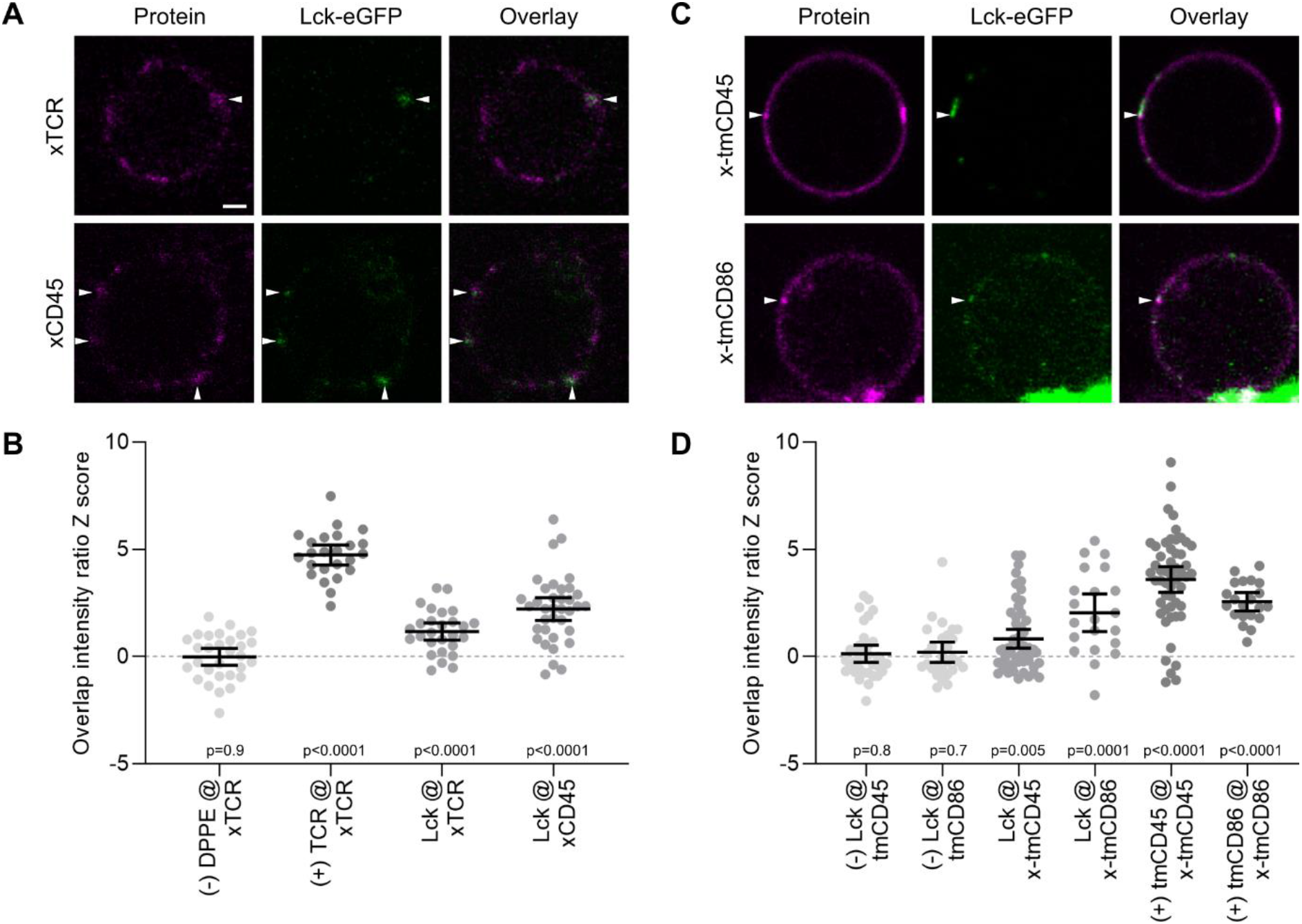
Co-localisation of Lck with protein aggregates in GPMVs. (A) Representative images of equatorial planes of GPMVs derived from Jurkat T cells expressing Lck-eGFP (green), with TCR (top) or CD45 (bottom) cross-linked with primary (OKT3 and Gap8.3, respectively) and fluorescently labelled secondary antibodies (magenta; scalebar: 2 μm). Arrows point to the sites of Lck co-localising with a protein aggregate. From the extracted two-colour intensity profiles along the membrane, (B) the overlap intensity ratio z-score was calculated (values >0 indicate above-random co-localisation; see Methods for details). As the positive and negative control (+/-), co-localisation of the TCR aggregates with TCR-mNeonGreen (TCR) and Atto647N-DPPE (DPPE) was investigated, respectively (images in Figure S8). Every data-point represents an intensity profile from a single image, the bars indicate means and their 95% confidence intervals. The p-values for each sample were calculated by a two-sided t-test against the hypothetical mean of 0 (all samples passed the normality tests). (C) As A, with transmembrane domains of CD45 and CD86 with fluorescent mRuby3, cross-linked with primary and fluorescent secondary antibodies. (D) As B, with non-aggregated proteins and self-co-localising aggregates as negative and positive controls (-/+). The p-values for each sample were calculated by the Wilcoxon sign-rank non-parametric test against the hypothetical median value of 0.

Interestingly, but perhaps not anymore unexpectedly, Lck was recruited also by aggregates of CD45 molecules (Figure 6A,B), which induced increased lipid ordering in GPMVs similarly as aggregates of TCR (Figure 3B). Moreover, even the aggregates of transmembrane domains of CD45 or CD86 were able to attract Lck (Figure 6C,D) as well as its membrane-binding domain SH4 (Figure S9). This diminishes the possibility of the observed co-localisation being the consequence of binding between intracellular protein domains, and invigorates the involvement of lipid-meditated interaction between the proteins upon aggregation. Whether this is primarily due to crowding or simply due to modulated diffusivity, as would arise upon ligand binding, remains to be investigated to ultimately discern the direct effects of the observed lipid-protein reorganisation in the context of T-cell activation.

### 2.6 Interplay of energy and entropy in lipid-protein interactions

The similarity in behaviour between aggregates of TCR and CD45, which both locally induce ordered lipid environments and attract Lo-associated Lck, as do cross-linked transmembrane domains of CD45 and unrelated CD86, hints towards a rather general biophysical mechanism at play. This notion can find broader support in numerous studies on lipid-protein interactions (*5*). For instance, simulations have shown that lipids in shells extending 3–4 nm around membrane proteins exhibit reduced mobility and increased order, which is further exacerbated by protein oligomerisation (*53*), immobilisation (*54*) or crowding (*55*), possibly resulting in nucleation of larger-sized ordered regions. It has indeed been shown experimentally that immobilisation of Lo-partitioning proteins can stabilise ordered membrane environments (*56*). Furthermore, cross-linking of lipids can change their phase partitioning towards more ordered environment (*57*), which can also be induced by immobilisation of certain lipid species (*45, 58*). In cells, the effect of such immobilisation can extend across to the opposing leaflet to assure co-registration of lipid ordering (*59*). This all indicates that protein oligomerisation, diffusion and lipid ordering are indeed tightly linked via passive biophysical concepts, and corroborates our findings that aggregation can modulate lipid-mediated interactions between proteins (Figure 6).

We finally aimed to roughly asses if this mechanism can apply also to partial immobilisation of a single molecule, as in ligand binding to the receptor. The observed 3–4-fold slow-down of TCR (Figure 5C) and therefore of its surrounding lipids corresponds to a reduction of their translational entropy (*60, 61*) by 1–1.5 *k*_*B*_ per molecule (Δ*S*_*t*_ = *k*_B_ ln(*D*_*slow*_/*D*_*free*_), with Δ*S*_*t*_ being the translational entropy change due to the change in mobility], *k*_B_ being the Boltzmann constant, and *D*_*free,slow*_ the diffusion coefficients of the freely moving and slowed-down molecules). To retain the equilibrium of the free energy (*F*) with lipid molecules in the shells of unrestrictedly moving proteins (*F*_*slow*_ = *F*_*free*_), the lipids around bound/aggregated proteins need to compensate this reduction of translational entropy by reducing their energy levels accordingly (Δ*E* = *T* Δ*S*, with Δ*E* being the reduction in energy due to the entropy reduction Δ*S*, with constant temperature *T*), achieved by stronger Van der Waals bonding between alkyl chains upon tighter packing (i.e., increased order). In fact, the corresponding energy decrease of Δ*E* = 1–1.5 *k*_B_*T* per molecule, further reinforced by reduced conformational entropy of lipids (*61, 62*) upon increased ordering, already exceeds the line tension-related demixing enthalpy needed to induce local phase separation (around 0.5–1 *k*_*B*_*T* per molecule) (*6, 63*). Taking into account that the shell of each protein contains tens of affected lipids, their collective contribution can overcome the energetic penalty of the protein in unfavourable lipid environment, which is on the order of 2 *k*_B_*T* per protein (*4*).

In this crude approximation, the observed degree of the slow-down of a membrane protein, which is compellingly comparable also to the reported relative difference in diffusivity between the Lo and Ld phases in model membranes (*63, 64*), can suffice to locally nucleate a more ordered lipid environment. Even as lipids dynamically exchange on a short time scale, these immobilisation-induced membrane patches represent an energetically favourable environment for membrane proteins that preferentially partition into ordered phases due to the length of their transmembrane domain and lipidation, such as Lck. Such attraction potentiates higher rates of their interaction with the ligand-bound membrane proteins than their freely moving copies. In the fair democracy of thermodynamics, the numerosity of lipid-protein and lipid-lipid interactions can thus provide a significant lever to orchestrate the nanoscale membrane organisation, which participate in membrane-hosted events such as signalling.

### 2.7 Further implications for cellular processes

Our data corroborate that membrane proteins can vary their preference for different lipid environments according to external changes. We have shown that membrane proteins that individually partition into disordered lipid environments, such as TCR or CD45, can upon aggregation shift their preference towards more ordered membrane parts. This could possibly be initiated already by partial immobilisation of the proteins upon ligand binding, as we attempted to demonstrate with slowing down the TCR by binding to lipids in supported lipid bilayers. Unfortunately, we could not directly follow the molecular events during the very early stage of T cell activation, as the required temporal resolution at single-molecule sensitivity is unattainable to the currently available live-cell microspectroscopy techniques. Furthermore, we conducted our measurements only 5–10 min after the cell-surface contact to avoid potential bias of the readouts owing to non-planar topography of the initial contact sites (*65*), during which cell activation could have induced additional aggregation of TCR. However, as simulations predict that the local lipid composition and order respond to immobilisation of a membrane protein within microseconds (*42*), this lipid-protein interplay may represent the very first subtle, though likely not decisive, membrane response in sensing of external binding.

We can speculate that the newly acquired tendency of the engaged TCR for more ordered lipid environment is further enforced by aggregation with the Lo-preferring co-receptor CD4 (Figure S4A), which additionally facilitates interactions with positive regulators Lck and LAT (Figure S4A), as we observed in our passive model system with GPMVs (Figure 6). The tightly packed membrane embedding also slows down the diffusion of the engaged protein complex through the close contact zone to promote phosphorylation (*66*). Furthermore, primary T cells with more ordered membrane, which is more prone to phase separation, have been found to form more stable immune synapses (*67*). The induced ordered regions can then further facilitate association with the cytoskeleton (*68*– *70*) needed for the translocation of the engaged and aggregated receptors for further processing and recycling.

Our findings therefore indicate that the key molecular events of the kinetic-segregation model of T-cell activation (*11*) could be accompanied by consistent changes in the membrane environment, supporting the sensitive and specific response of each T cell. However, from the presented data we cannot attest whether the seen effects can in any way assist the activation, or they arise as its side effect. It therefore remains to be quantified whether the observed membrane remodelling affects the frequency of interactions between the investigated proteins, as well as with other negative regulators, such as Lck-inactivating protein Csk (PAG) (*4*) and SHP-1 (*71*), which also partition into ordered membrane environments. This will help to determine the degree to which the suggested lipid-mediated binding recognition upon receptors’ immobilisation can influence the final signalling outcome, and perhaps its relation to malfunctions such as immune disorders.

Similarities of molecular machinery employed in other cell types will enable a direct translation of these findings to other signalling events, in particular to triggering of B lymphocytes, which involves phosphorylation of B-cell receptors (BCR) controlled by a kinase Lyn and phosphatase CD45. Remarkably, it has recently been shown that during this process, aggregation of the BCR, itself preferring disordered membrane environment, promotes accumulation of more ordered lipid regions, recruitment of thereto-partitioning Lyn and exclusion of CD45, which is largely powered by lipid-lipid interactions for soluble antigen (*31*) and additionally enhanced for surface-bound antigens (*32*). Similar parallels could be drawn also to the activation of other cascades that knowingly depend on lipid heterogeneities, such as FcεRI-mediated IgE signalling in mast cells (*33, 34*).

3 Acknowledgements

The authors greatly thank prof. Simon J. Davis for stimulating discussions, drs. Christoffer Lagerholm, Silvia Galiani, Jana Koth, and Dominic Waithe from the Wolfson Imaging Centre Oxford for their support with imaging infrastructure, Drs. Katharina Reglinski and Dilip Shrestha for their help with lab work, and Andrey Klymchenko for kindly providing the probe NR12S. The work was funded by the European Commission via Marie Skłodowska-Curie individual fellowships to I.U. (grant no. 707348) and E.S. (627088), the Wolfson Foundation (No. 18272), the Medical Research Council (MRC, grant no. MC_UU_12010/unit programs G0902418 and MC_UU_12025), MRC/BBSRC/EPSRC (grant no. MR/K01577X/1), the Wellcome Trust (grant no. 104924/14/Z/14 to C.E. and 098274/Z/12/Z to S.J.D.), the Deutsche Forschungsgemeinschaft (Research unit 1905 ‘Structure and function of the peroxisomal translocon’, Jena Excellence Cluster “Balance of the Microverse”, Collaborative Research Center 1278 ‘PolyTarget’), the Jena Center of Soft Matter, and Oxford internal funds (John Fell Fund and EPA Cephalosporin Fund), and Balliol College Oxford. E.S. was funded by SciLifeLab Fellowship and Karolinska Institutet Foundation grants.

## 4 Methods

### 4.1 Plasmids

For transient transfections, a pHR plasmid backbone was modified to contain a BamHI-MluI-SpeI-NotI multiple cloning site. A Kozak sequence, signal peptide, HA tag and GGSG linker were added between BamHI and MluI. Full length CD28 (aa 19-220, UniProtKB P10747) and CTLA-4 (aa 36-223, UniProt P16410) were cloned between MluI and SpeI. Similarly, the transmembrane (tm) domains of human CD4 (aa 391-423 UniProtKB P01730), CD45 (aa 573–608, UniProtKB P08575), CD86 (aa 243-277, UniProtKB P42081) and LAT (aa 1-35, UniProtKB P43561), and their mutants, were cloned between MluI and SpeI, and for all constructs fluorescent protein mNeonGreen (mNG) or mRuby3 between SpeI and NotI. Lck (SH4 domain) and mutants were cloned between BamHI and SpeI with only a Kozak sequence. For stable cell lines, using a pHR plasmid backbone, mNG was fused to the C terminus of the beta chain of a high affinity gp100 specific TCR (kind gift from Immunocore Ltd.), and eGFP to the C-terminus of wild type human Lck (UniProtKB P06239).

### 4.2 Cell culturing and transfections

Adherent HEK 293T cells (CRL-3216, ATCC), and derivatives, were grown in T75 flasks (CELLSTAR) with DMEM (D5796, Sigma-Aldrich) supplemented with 10 % foetal bovine serum (FBS; Gibco), 2 mM L-glutamine and 1 % penicillin-streptomycin (all from Sigma-Aldrich). Jurkat T cells (ATCC), and derivatives, were cultured in T25 flasks (CELLSTAR) with RPMI (1640, Thermofisher) supplemented with 10 % FBS (Gibco), 10 mM HEPES, 2 mM L-glutamine and 1 % penicillin-streptomycin (all from Sigma-Aldrich). HEK 293T and Jurkat cells were grown at 37 °C and 5 % CO_2_. Suspension Chinese hamster ovary cells (FreeStyle CHO-S cells, R80007, Thermofisher) were kept in suspension in plastic Erlenmeyer shake flasks (Corning) with FreeStyle CHO Expression Medium (12651022, Thermofisher) supplemented with 8 mM L-glutamine (Sigma-Aldrich) under 37 °C, 85% humidity, and 8 % CO_2_ on an orbital shaker at 125 rpm. All cells were free of mycoplasma.

Lentiviral production and transduction were used to generate stable Jurkat cell lines expressing gp100-reactive TCR-mNG, Lck-eGFP or Lck (SH4)-mNG, and CHO-S cells expressing non-fluorescent TCR, additionally assured with fluorescence-assisted cell sorting. For transient transfection of membrane proteins, HEK 293T cells were first seeded into 6-well plates, coated with poly-L-lysine (Sigma-Aldrich), one day prior to transient transfection using 0.5 μg of DNA per well and GeneJuice (Merck Millipore) as per the manufacturer’s instructions, and left incubating for another 2–3 days before GPMV preparation.

### 4.3 Preparation of giant plasma-membrane vesicles (GPMVs)

For GPMVs prepared from activated T cells, these were first incubated in HEPES buffered saline (HBS; 10 mM HEPES, 150 mM NaCl; pH 7.4) with 10 μg/ml OKT3 anti-CD3 antibody (kindly provided by the Human Immunology Unit, WIMM) for 3 min at room temperature, followed by the GPMV formation procedure, described below.

GPMVs from adherent HEK 293T cells or Jurkat and CHO cells in suspension, were prepared following the established protocol (*35*) with necessary modifications for suspension cells (*50*). Cells were first pre-swelled by washing in 30 % GPMV buffer (10 mM HEPES, 150 mM NaCl, 2 mM CaCl_2_; pH 7.4), followed by a 2-hour incubation at 37 °C in full GPMV buffer with n-ethylmaleimide (NEM, final concentration 2 mM), or dithiothreitol (DTT, 25 mM) and paraformaldehyde (PFA, 0.07 %). For this step, suspension cells were transferred into 35-mm cell-culturing plastic petri-dishes (Cornig). The harvested GPMVs were then purified by spinning down the cellular debris (2 min at 1 krpm) and concentrated by further centrifugation (15 min at 10 krpm).

For subsequent aggregation of proteins in GPMVs, the vesicles were incubated at room temperature with approx. 0.1 μg/ml of the corresponding primary antibody (OKT3 anti-CD3 and Gap8.3 anti-CD45, kindly provided by Davis lab (*72*); both mouse anti-human) for 3–10 hours, followed by another 2–6 h of incubation with approx. 0.3 μg/ml of a secondary goat anti-mouse antibody – bare or labelled with Alexa Fluor 488, 568, or 633 (Thermofisher). Transmembrane protein constructs with extracellular HA-group were cross-linked with Biotin anti-HA.11 Tag Antibody (BioLegends) and Streptavidin Alexa Fluor 647 (Thermofisher).

### 4.4 Antigen-binding fragments and proteins for SLB decorations

cDNA encoding extracellular domain (ECD) of CD58 (residues 29–215, UniProtKB P19265), the ECD of HLA-A alpha chain (residues 25-269, UniProtKB B1PKZ1) and beta-2-microglobulin (β2M, residues 21-119, UniProtKB P61769) were ligated into pHR downstream of the sequence encoding a Kozak sequence (GCCACC) and cRPTPs signal peptide. For the UCHT-1-CD45R0-C-tag/His-tag fabs, the UCHT-1 variable domain sequences were obtained from a previous report (*73*). Both the V_H_ and V_L_ domains were cloned into their respective pOPINVH and pOPINVL backbones. Only pOPINVH was modified to have the large protein CD45RO fused to the C-terminus of the V_H_ constant region with either His tag or C-tag (EDQVDPRLIDGK) at the C terminus of CD45R0. For CD58, HLA-A alpha chain and UCHT-1CD45R0-His-tag sequences a H6-SRAWRHPQFGG-H_6_ ‘double His_6_’ tag was added on the C-terminus. Soluble pMHC (HLA-A and β2M), CD58 and UCHT-1-CD45R0-C-tag/His-tag were produced as previously described (*52, 74*) (Chen et al., submitted). Briefly, the ECD of CD58 was expressed transiently in HEK 293T cells, whereas the alpha and beta chains for pMHC (HLA-A) were purified from inclusion bodies and folded in the presence of gp100 (agonist; YLEPGPVTV; GenScript). UCHT-1VH-CD45R0-C-tag/His tag vector was co-expressed along with UCHT-1VL transiently in HEK 293T cells. Proteins were either purified using NI-NTA agarose (Qiagen) or anti C-tag beads. Monomers were isolated by fast protein liquid chromatography using an AKTA pure protein purification system.

### 4.5 Preparation of antibody-coated glass surfaces and supported lipid bilayers (SLBs)

For immobilisation of proteins in the membrane of live cells, glass-bottom 8-well microscopy chambers (ibidi μslides, glass thickness no. 1.5) were incubated with 1 μg/ml of OKT3 or Gap8.3 and 9 μg/ml of OX7 (non-specific anti-rat Thy1.1) in HBS overnight at 4 °C, as applied before (*28*). Prior to use, chambers were washed at least three times with HBS.

Glass-supported lipid bilayers (SLBs) were prepared using vesicle fusion (*75*). Lipid mixture consisting of 98% POPC (Avanti Polar Lipids) and 2% DGS-NTA-Ni^2+^ (Ni; Avanti Polar Lipids) in chloroform were mixed in a round bottom glass flask and dried under a stream of nitrogen. The dried lipid mix was resuspended in 0.22-μm filtered PBS, vortexed, and tip sonicated for 30 minutes on ice to produce small unilamellar vesicles (SUVs). Glass coverslips (25 mm, thickness no. 1.5; VWR) were cleaned overnight in 3:1 sulfuric acid/hydrogen peroxide at room temperature, rinsed in MQ water, and plasma cleaned for 1 minute. CultureWell 50-well silicon covers (Grace Bio-Labs) were cut and placed on the washed coverslips (max 4/coverslip). SUVs were added to each well at a final concentration of 0.5 mg/ml (10 μl) and left for 0.5–1 hours at room temperature. Wells were washed by removing and adding 5 μl PBS x5 before adding proteins at the desired density. Double His_6_ were used to tether the proteins to the lipid bilayer via interaction with DGS-NTA-Ni^2+^, providing more stable binding to SLBs than single His_6_ tagged proteins (*76*). Final concentration of protein mixes included 1 ng/μl pMHC-gp100, or 1 ng/ul CD58 and 1 ng/μl pMHC-gp100. Protein mixes were incubated with the bilayer for an hour at room temperature and washed in pre-warmed PBS (37°C) five times immediately before use. Protein densities were matched to physiological levels: pMHC-gp100 (agonist) 50–100/µm^2^ and CD58 300–500/µm^2^. Large densities of non-agonising/non-binding double His_6_ pMHC were used to block unbound nickel sites. Fluorescence correlation spectroscopy (FCS) was used to relate protein concentration to density on the SLB (data not shown).

### 4.6 Two-colour imaging of proteins in phase-separated GPMVs

For experiments with phase-separated GPMVs, the membranes were labelled with a fluorescent lipid analogue Atto647N-DPPE (AttoTec) by incubating the GPMV solution with the dye at up to 0.02 μg/ml for 5–10 min, followed by a 20-min incubation with up to 0.2 mM deoxycholic acid (DCA) to facilitate large-scale phase separation (*46*). A drop of the GPMV suspension was then transferred into the observation chamber created with Vaseline between two coverslips (22 × 50 mm, thickness no. 1.5; by Menzel), which was adhered to a Peltier cooling device set to 4 °C and mounted on the microscope stage (*35*).

The observations were conducted using an inverted laser-scanning confocal microscope Zeiss LSM780 equipped with a 40x/1.1 water-immersion objective. Green and far-red fluorescence was excited with lasers at 488 nm and 633 nm and collected in two channels within the spectral windows of 500– 590 nm and 640–700 nm. For each vesicle, a z-stack of images across the entire vesicle was collected with spacing between slices of 0.56 μm and sampling of around 0.1 μm/pixel. Acquisition of each slice took 2–6 seconds, which was fast enough that the locations of protein aggregates were not smeared by their diffusion.

### 4.7 Analysis of protein partitioning in phase-separated GPMVs

The preference of membrane proteins for the characteristic lipid phases in phase-separated GPMVs was analysed using Fiji/ImageJ as described before (*35*). In brief: along a manually defined line connecting regions with ordered and disordered membrane phases on the opposite sides of the vesicle (Figure S1A), a profile of the protein’s intensity was extracted (Figure S1B). The peak intensity values in the liquid ordered and disordered phase (*I*_*Lo*_ and *I*_*Ld*_, respectively) were read out and used to calculate the Lo-partitioning percentage (P_Lo_):

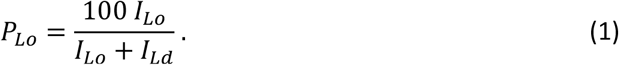

For the proteins with frequent partitioning at the border between the two membrane phases, two-colour intensity line profiles along the membrane of each vesicle were manually extracted using Fiji/ImageJ (Segmented Line tool with line width of 5 pixels and Spline fit enabled) and further analysed with custom scripts written in Mathematica. The profiles were first de-noised using a Wiener filter, pertaining the peak features. Proteins and lipid-phase boundaries were identified in the green channel (protein) and gradient-filtered red channel (Atto647N-DPPE), respectively, using the PeakDetect function. For each identified border (170–300 borders from 50–70 intensity profiles per sample), the distance to the closest protein was recorded (*d*_*min*_, as indicated in Figure 1C and exemplified in Figure S3C). For all borders from each sample, the cumulative distribution function of minimal border-to-protein distances was calculated (*CDF*(*d*_*min*_), Figure S3D), and compared to those generated from the same profiles with the two channels slid with respect to each other for a random spatial shift (*CDF*_*rand*_(*d*_*min*_)):

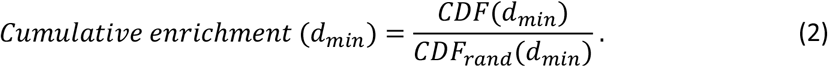

Figure 1C and Figure 2D display the averages and standard deviations of the *Cumulative enrichment* factor calculated for 200 randomly shifted profiles per analysed image; values above 1 thus indicate above-random occurrences of *d*_*min*_.

### 4.8 Spectral imaging of GPMVs and live cells

To monitor membrane order in GPMVs, these were incubated with the polarity-sensitive membrane dye NR12S (*47*) at a concentration of 0.05 μg/ml for 30 min. Homogeneous vesicles were imaged at room temperature in plastic-bottom 8-well microscopy chambers (ibidi μslides), whereas for phase separation at 4 °C, Vaseline-sealed observation chambers were used, as described above.

To measure membrane order in live cells with respect to a protein’s location, cells were first incubated in HBS with the corresponding antigen-binding fragment (fab): plain Jurkats with anti-CD3 or -CD45 fabs with Alexa Fluor 488 at 0.2 μg/ml on ice, Jurkat TCR-mNG with 50 ng/µl of UCHT-1CD45RO-C-tag/His-tag fab in RPMI without supplements at 37 °C. After 15 min, NR12S was added (0.5 μg/ml), and cells were left to incubate for another 5 min before washing with warm HBS. Cells were then carefully dropped onto the surface of interest (antibody-coated coverslip or protein-decorated SLB) on the microscope stage (37 °C and 5 % CO_2_) and left for 5–10 min to settle and spread before imaging.

Spectral imaging was conducted using the same microscope as above (or a newer model Zeiss LSM880 with equivalent functionality). Z-stacks of GPMVs were taken with a 40x/1.1 water-immersion objective, as described above, whereas the bottom membranes of live cells were imaged with a 63x/1.4 oil immersion objective with spatial sampling around 90 nm/pixel. Fluorescence was excited at 488 nm and detected in the so-called “Lambda Mode” with 22 channels within the spectral range 500–695 nm, i.e. with a wavelength step of 8.9 nm.

### 4.9 Analysis of spectral shifts in GPMVs and live cells

To decompose the spectrum in every pixel of the image into the contributions from the protein (eGFP, mNeonGreen, or antibody with Alexa Fluor 488) and the polarity-sensitive membrane probe NR12S, both excited with the 488-nm laser, spectral fitting-based (*49*) linear-unmixing was employed using custom software written in Mathematica (available upon request). Each spectral line-shape was approximated with a transformed log-normal distribution (*49*), described by three parameters: spectral peak position (λ_MAX_), full-width at half-maximum (*w*), and asymmetry (*a*). Except from λ_MAX_ of NR12S and the relative intensity ratio of the two fluorophores, all other spectral parameters were fixed to values optimised on measurements with individual fluorophore. Prior to fitting, the spectra of 3×3 neighbouring pixels were moving-averaged to improve the signal-to-noise ratio and thereby increase spectral sensitivity, and background spectrum from a neighbouring region subtracted.

To determine the spectral shifts of NR12S at protein aggregates in GPMVs (Figure 3 and Figure S5), aggregates were identified by intensity thresholding, and average peak position of NR12S within the masked region (λ_MAX,*at protein*_) compared to membrane parts without protein aggregates (λ_MAX,*elsewhere*_):

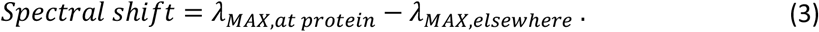

In live cells (Figure 4 and 5), due to less pronounced protein aggregates and higher variations of the NR12S intensity, pixels were classified into groups with high and low local protein concentration based on the relative intensity of the protein signal relative to the membrane intensity (*I*_*protein*_/*I*_*NR12S*_), with thresholding at the median value within each cell. Only central regions of the cells (i.e., with only the bottom membrane in the focus) were analysed. Pixels with a low portion of the membrane signal, where determination of λ_MAX_ was potentially influenced by miss-fitting of the tails of the protein spectrum, were also deemed unreliable and therefore discarded.

To ensure comparability and avoid bias, the routines were fully automatized and repeated with the same set of parameters across all comparable samples. Statistical analysis was performed with GraphPad Prism, as described in each figure caption.

### 4.10. Diffusion measurements of proteins in live cells

The diffusion coefficient of TCR-mNG in the bottom membrane of Jurkat T cells spreading on SLBs (prepared as for spectral imaging), was measured by fluorescence correlation spectroscopy (FCS) using a customised setup (*77*) built around a RESOLFT microscope (Abberior Instruments), controlled via Imspector software, and equipped with a 100x/1.4 oil-immersion objective (Olympus) and a stage-top incubator (Okolab) set to 37 °C. The sample was excited with a 488-nm diode laser (PicoQuant) pulsing at 80 MHz and average power 2 μW, measured at the back aperture of the objective, and fluorescence collected by an avalanche photodiode (APD, Excelitas) within the spectral window of 510–565 nm. The signal from the detector was correlated with a hardware correlator (correlator.com Flex02-08D) with typical acquisition times of 10–15 s. Acquisition sites were manually chosen in membrane areas with high protein density.

The obtained FCS curves were analysed with FoCuS-point fitting software (*78*) (freely available at https://github.com/dwaithe/FCS_point_correlator), using the standard model of anomalous 2D diffusion with two triplet components (*79*) with their correlation times fixed to 0.1 and 2 ms (Figure S7). From the extracted transit times (*τ*_xy_) and estimated diameter of the effective confocal observation spot (*d* = 200 nm, full-width at half-maximum), the diffusion coefficient (*D*) was calculated using (*80*)

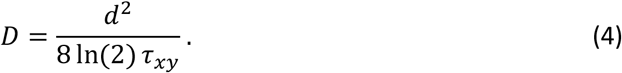

For the activating condition (adh+pMHC in Figure 5C), it was challenging to obtain converging and thus reliable correlation curves, for which reason the number of measured instances is lower.

### 4.11 Co-localisation of proteins in GPMVs

For two-colour co-localisation experiments, GPMVs with antibody-stained protein aggregates (or with Atto647N-DPPE in the negative control, as above) were imaged in plastic-bottom 8-well microscopy chambers (ibidi μslides) at room temperature, using the same instrumentation and settings as for observations of phase-separated GPMVs, described above. The red fluorescence of transmembrane proteins with mRuby3 was excited at 561 nm and collected within 580–630 nm, in a line-interleaved mode with the green channel.

To quantify the co-localisation of protein aggregates in two-channel images of GPMVs, intensity line profiles along the membrane of each vesicle were manually extracted using Fiji/ImageJ (Segmented Line tool with line width of 5 pixels and Spline fit enabled) and further analysed with custom scripts written in Mathematica. The profiles were first de-noised using a Wiener filter, pertaining the peak features. By thresholding the data in the reference channel (e.g. of TCR aggregates) at an intensity value at a predefined factor above the background, a mask was created to extract the average intensities of the other channel (e.g. Lck-eGFP) within and outside of the masked regions (*I*_*at protein*_ and *I*_*elsewhere*_, respectively, see Figure S8B), used to calculate the *Overlap Intensity Ratio* (*OIR*):

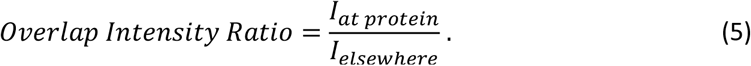

To compare this to a random distribution, *OIR* was calculated for the intensity profile of the channel of interest split in 50-pixel sections and randomly reshuffled 1000 times (*RandomOIR*), from which the mean and standard deviation were calculated to yield a *Z score* of the experimental *OIR*:

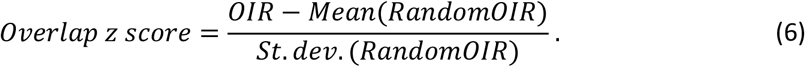

Hence, positive values of the Overlap z score indicate above-random co-localisation of signals in the two channels. Statistical analysis was performed with GraphPad Prism, as described in each figure caption.

## 6 Supporting Information

### 6.1 Partitioning of proteins in phase-separated GPMVs

**Figure S1:**
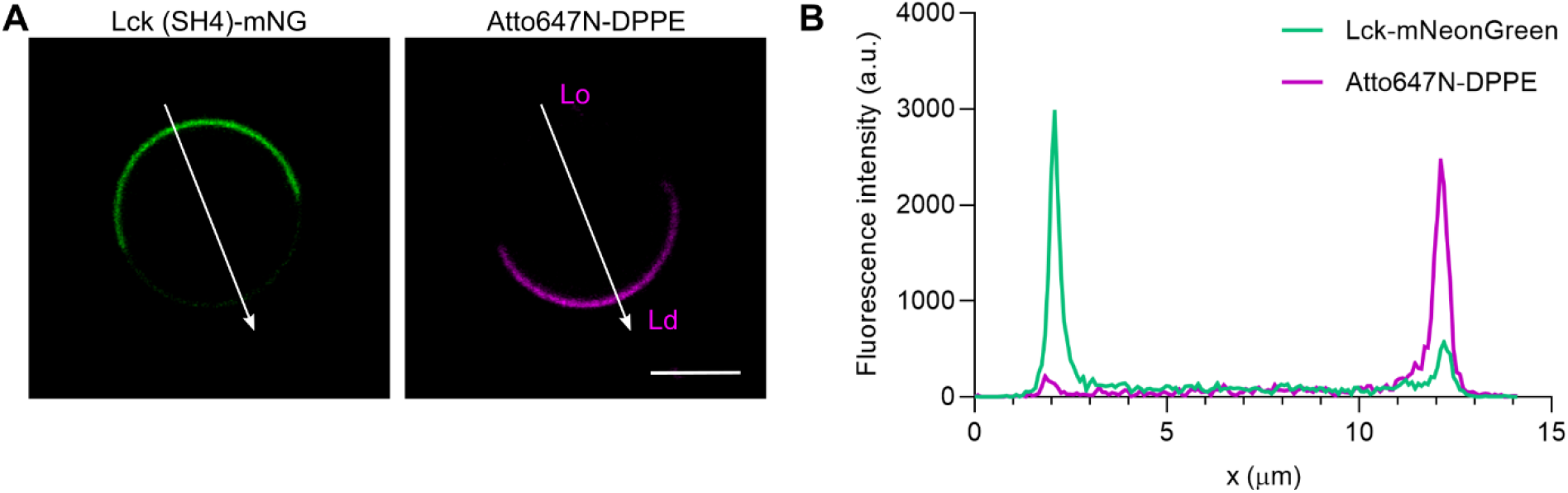
Determination of proteins’ Lo partitioning percentage. (A) In an image of the equatorial plane of a GPMV (scale bar: 5 μm) with the protein of interest (here Lck SH4-mNeonGreen in green, left) and the liquid-disordered phase marker Atto647N-DPPE (magenta, right), a manually-selected line connecting opposing membrane regions with ordered (Lo) and disordered (Ld) lipid phase was used to extract (B) the intensity profile of the protein (green). The peak intensity values at the membrane were used to calculate the Lo partitioning percentage (see Methods).

**Figure S2:**
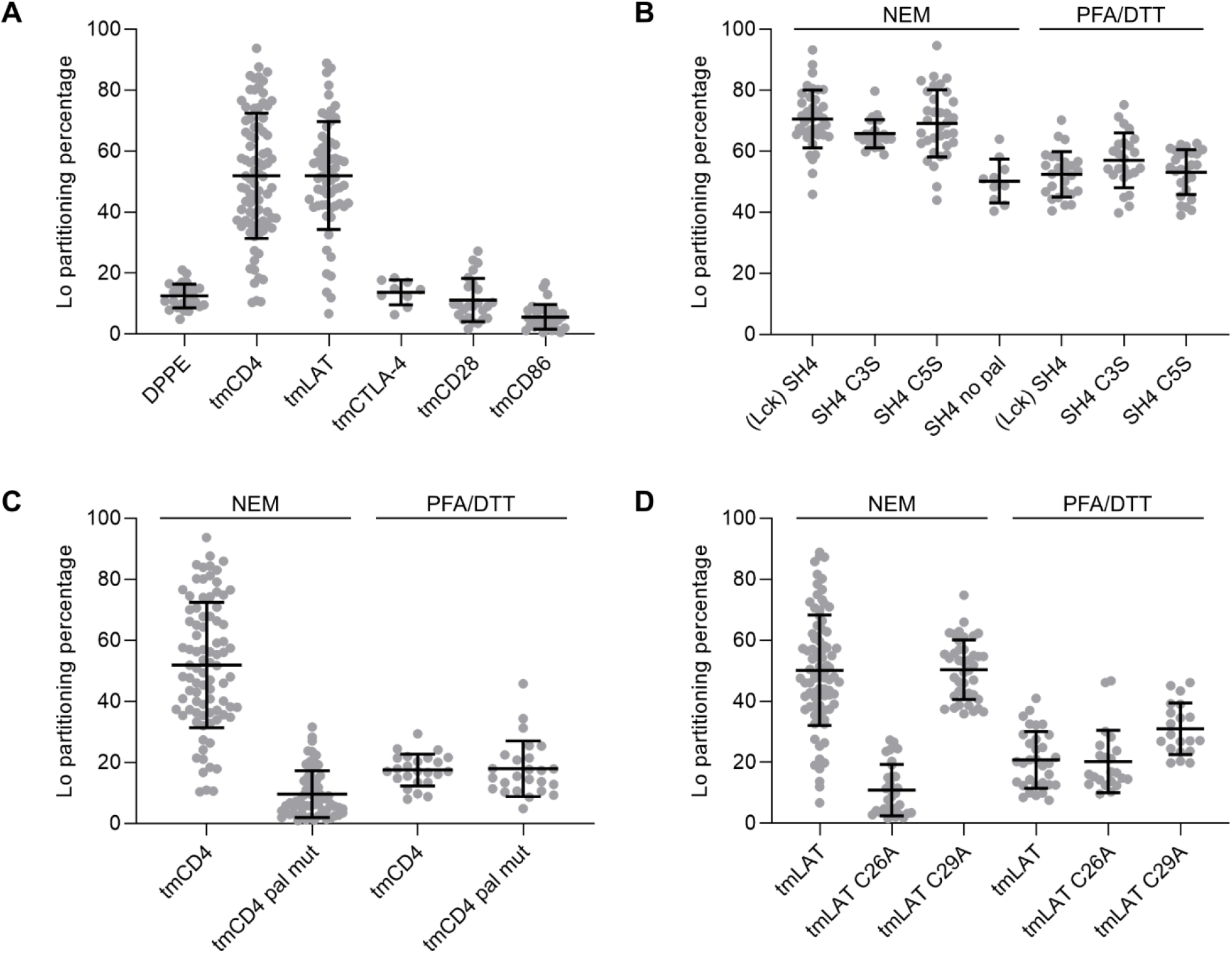
Lo partitioning percentages for transmembrane (tm-) protein domains and their membrane-anchor mutants in GPMVs prepared from HEK 293T cells with different procedures (NEM or PFA/DTT): (A) wildtype proteins, (B) Lck, (C) CD4, and (D) LAT. Every data-point represents a value from a single GPMV, the bars indicate the means and standard deviations.

**Figure S3:**
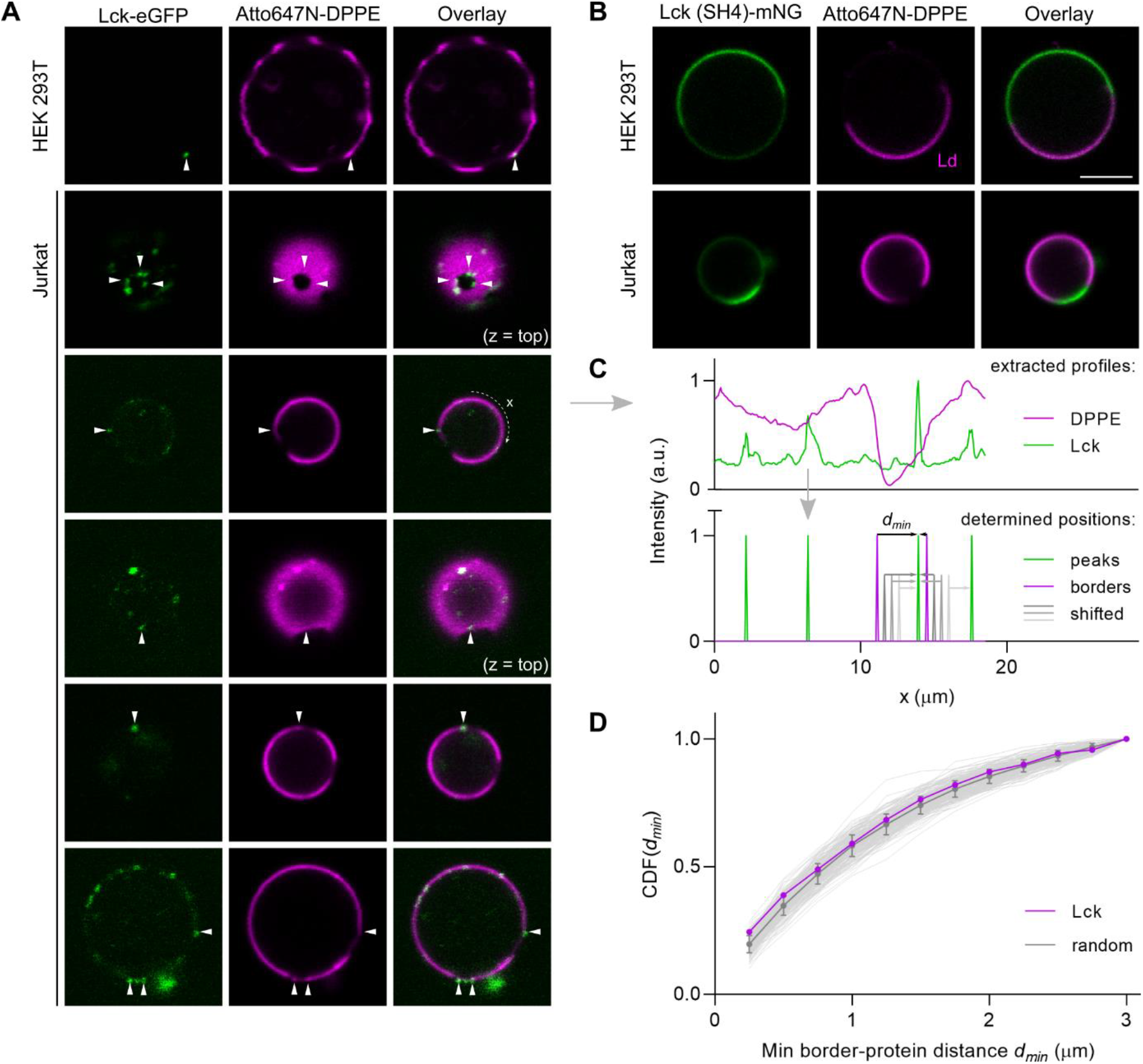
Partitioning of fluorescent Lck constructs in GPMVs. Representative images of phase-separated GPMVs from HEK 293T and Jurkat T cells, expressing (A) full-length Lck, fused to eGFP (green), or (B) its membrane-binding domain SH4 tagged with mNeonGreen. The vesicles were stained with the liquid-disordered (Ld) phase marker Atto647N-DPPE (magenta; scale bar: 5 μm). Two images in (A) were taken in the plane close to the top of the vesicle, as indicated (z = top). Arrows point at proteins localised at the border of lipid phases. (C) To evaluate the border partitioning of proteins or their clusters, two-colour intensity profiles were extracted (top), in which the lipid-phase borders and protein clusters were identified (bottom). For each border, the distance to the closest protein (*d*_*min*_) was calculated. (D) From all images of each sample, the cumulative distribution function (CDF) of *d*_*min*_ was calculated (magenta) and compared to those obtained with randomly shifted intensity profiles (grey, as indicated in panel C), to obtain the Cumulative enrichment plots (Figure 1C, Figure 2D; see Methods for details).

### 6.2 Partitioning of protein clusters in phase-separated GPMVs

**Figure S4:**
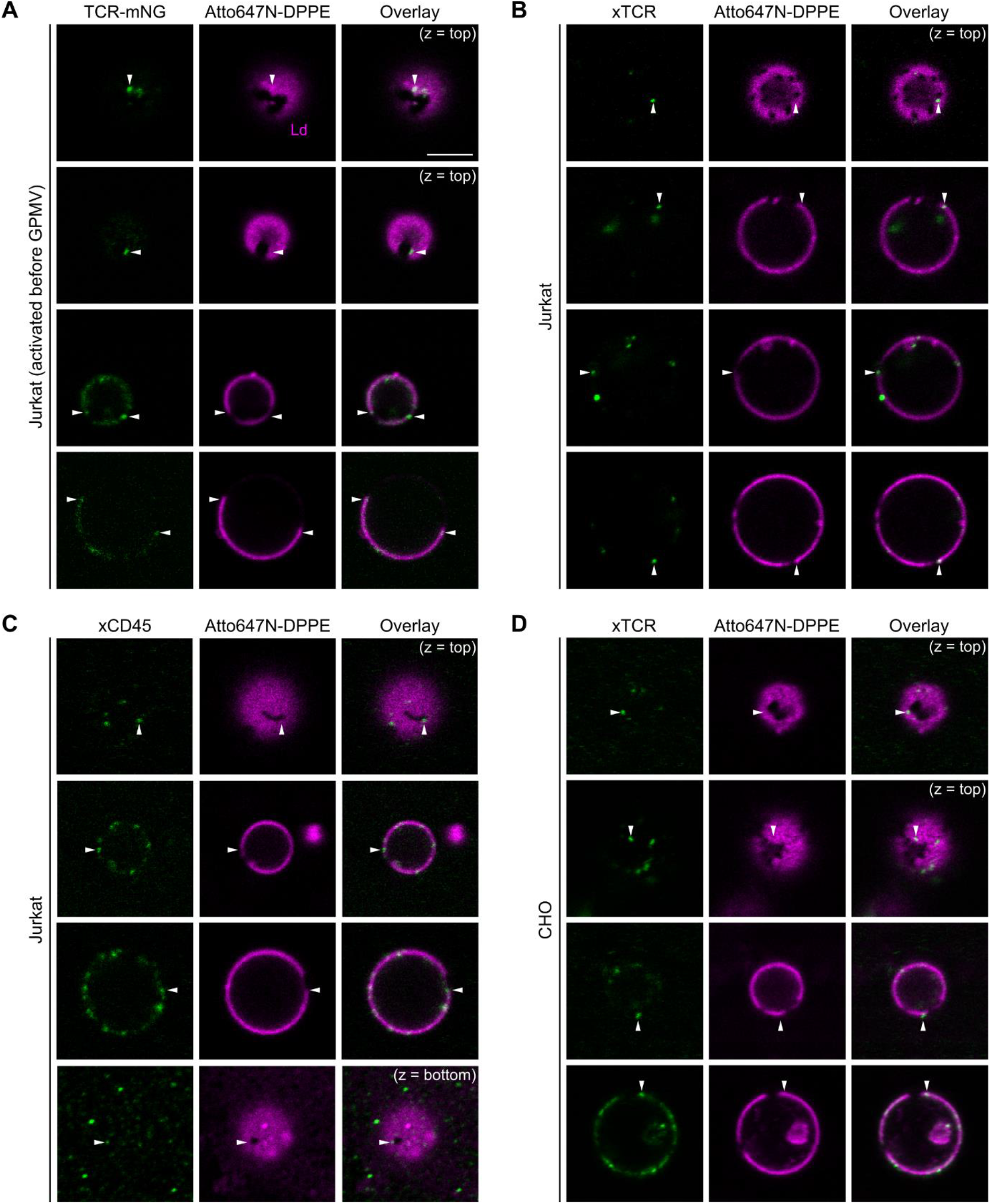
Partitioning of protein clusters (green) and the Ld phase marker Atto647N-DPPE (magenta) in phase-separated GPMVs. Representative images of (A) TCR-mNeonGreen in GPMVs from Jurkat T cells pre-activated by OKT3 in solution; (B) TCR and (C) CD45 from Jurkat T cells, and (D) TCR from CHO cells, all of which were cross-linked in GPMVs by primary and fluorescent secondary antibodies. Arrows point at proteins localised at the border of lipid phases. Other-than-equatorial imaging plains (z) are indicated.

### 6.3 Membrane order in GPMVs

**Figure S5:**
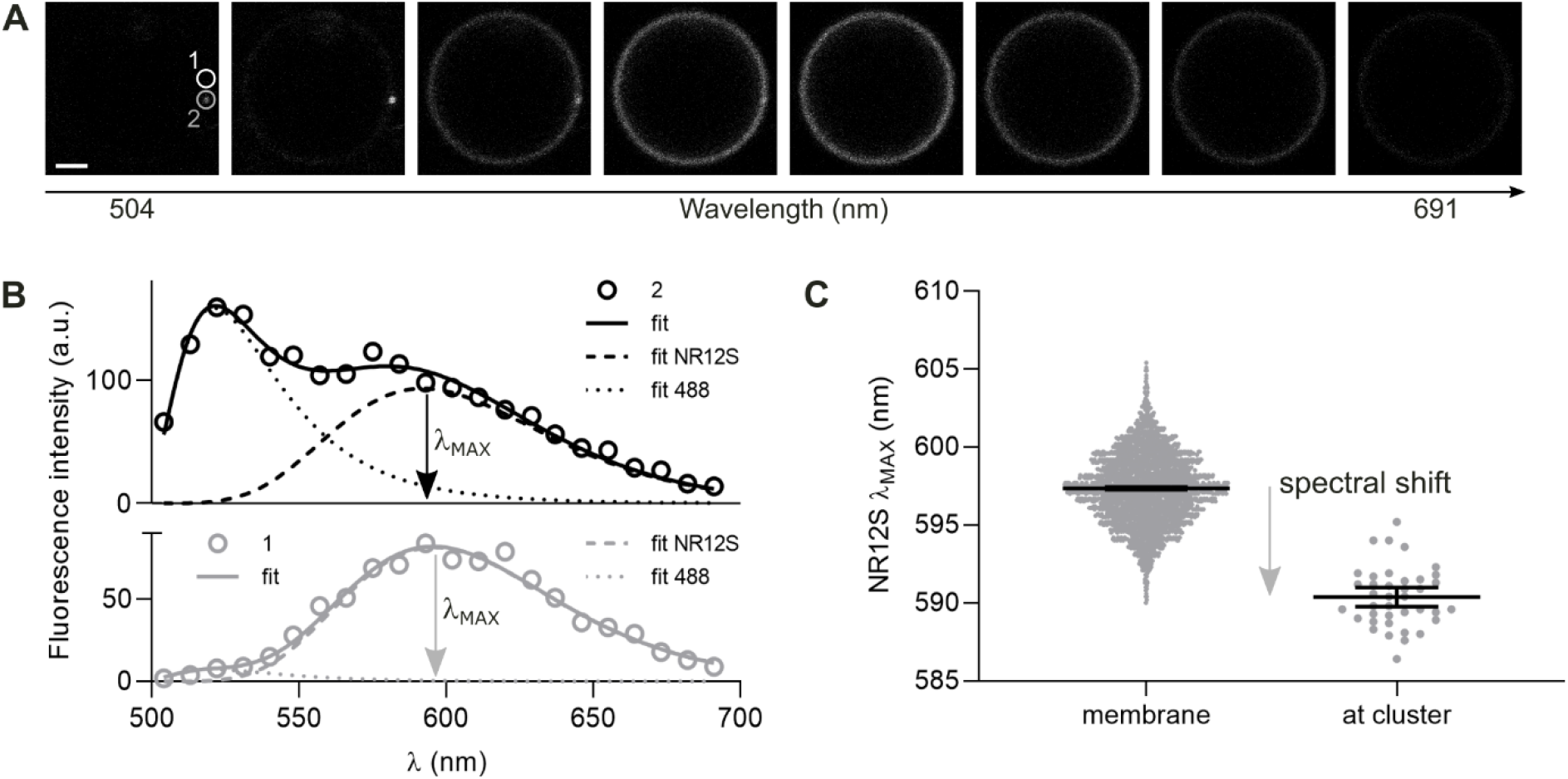
Determination of membrane order in GPMVs with proteins clustered by antibodies. (A) Spectral series of images of an exemplary GPMV with TCR clustered with 488-labelled antibodies and stained with the polarity-sensitive membrane dye NR12S, excited simultaneously by a blue laser (scale bar: 2 μm). The encircled regions indicate locations from which we extracted (B) exemplary fluorescence emission spectra (correspondingly labelled as 1 and 2). Each spectrum was fit (solid lines) by two spectral components – for NR12S and the protein-bound 488 antibody (dashed and dotted lines, respectively). The fit parameters from each pixel, i.e. fitted intensities of the 488 antibody and NR12S as well as its spectral peak position (λ_MAX_), were used to generate the spatial maps (Figure 3A). Intensity values of 488 were used to identify protein clusters. (C) Per-pixel values of λ_MAX_ at the site of each protein cluster were compared to those from the rest of the membrane, and the difference between their means was used as the spectral shift of NR12S for each identified protein cluster, displayed in Figure 3B. Bars indicate means and their 95% confidence intervals.

**Figure S6:**
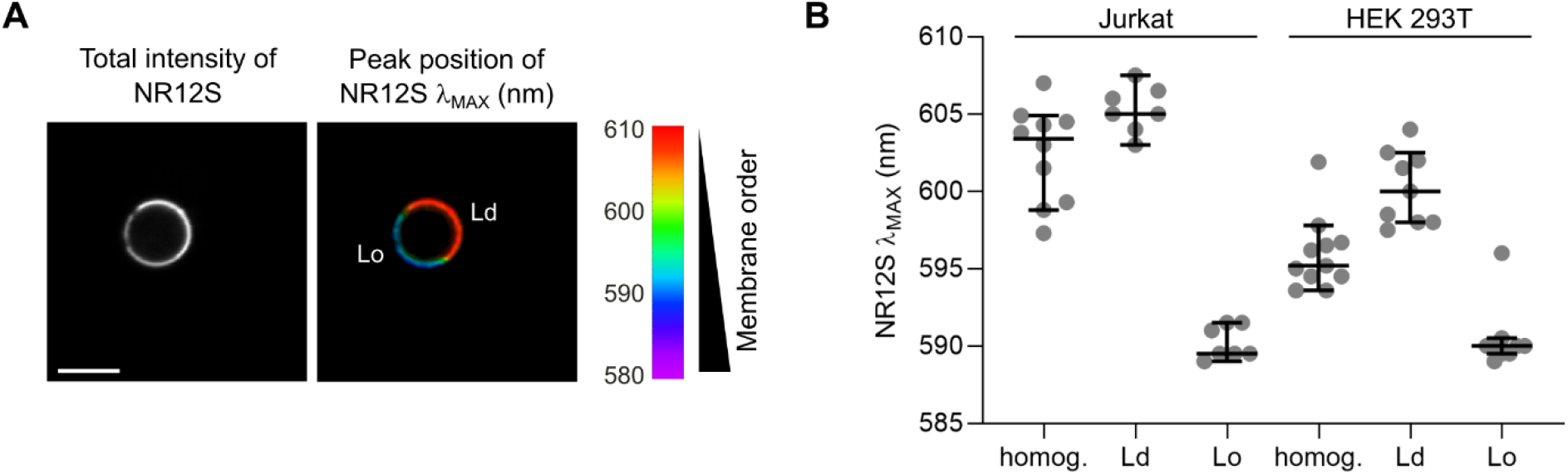
Membrane order in phase-separated GPMVs. (A) Total intensity of NR12S (left) and its fitted peak position (λ_MAX_, right) for an exemplary GPMV (scale bar: 5 μm). (B) Peak positions of NR12S in individual homogeneous GPMVs or in both phases of separated vesicles (disordered Ld, ordered Lo), derived from Jurkat and HEK 293T cells. Bars indicate medians and their 95% confidence intervals.

### 6.4 Diffusion measurements of TCR in T cells on SLBs

**Figure S7:**
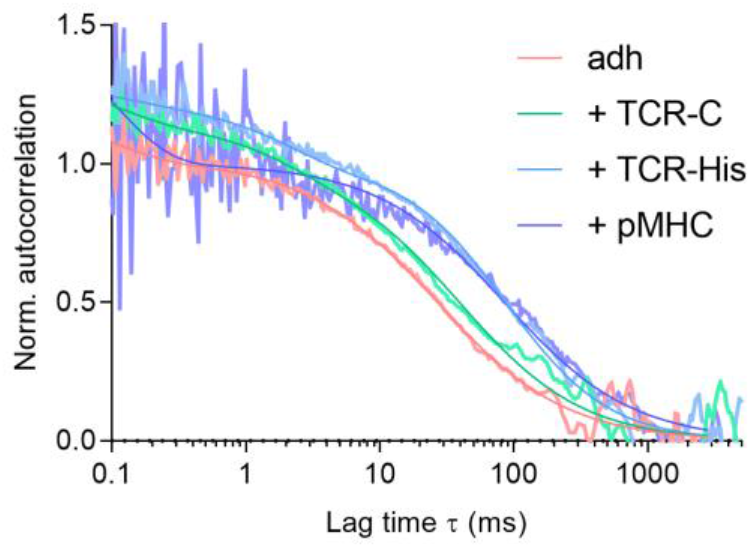
Exemplary normalised FCS curves, and fits thereto (smooth lines of corresponding colour), of TCR-mNeonGreen in the bottom membrane of Jurkat T cells spreading on SLBs under different conditions, as described in Figure 5.

### 6.5 Co-localisation of Lck with protein clusters

**Figure S8:**
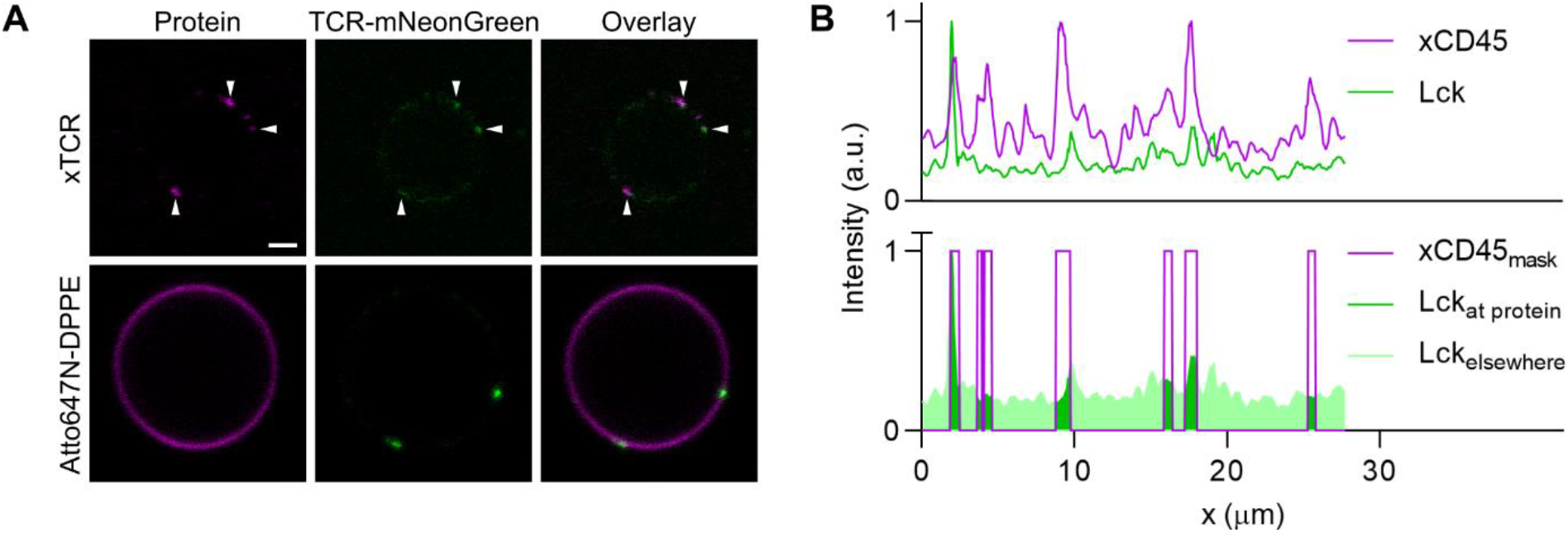
Controls for co-localisation with protein clusters in GPMVs, derived from Jurkat T cells expressing TCR-mNeonGreen (green). TCR was clustered with primary (OKT3) and fluorescent (top, magenta) or non-fluorescent (bottom) secondary antibodies. The latter were labelled with a fluorescent lipid analogue Atto647N-DPPE (magenta). Scalebar: 2 μm. (B) Exemplary intensity profiles (top) along the perimeter of a GPMV with Lck-eGFP and cross-linked CD45. Thresholded values of the latter were used as a mask (bottom) to determine the average Lck intensity at the sites of protein clusters and elsewhere, used in the calculation of the Overlap Intensity Ratio (see Methods for details).

**Figure S9:**
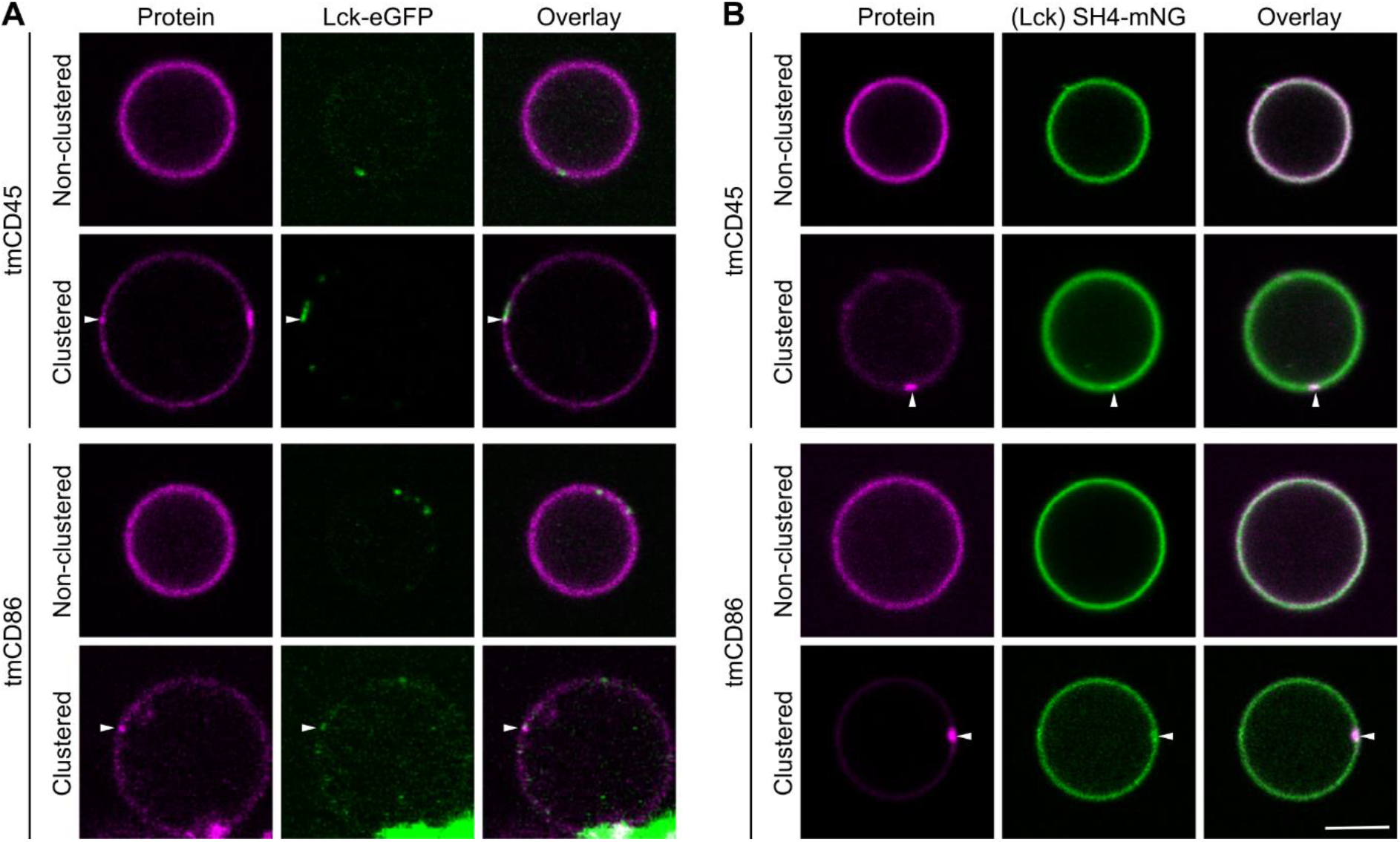
Co-localisation of fluorescent protein constructs in GPMVs from Jurkat T cells. Representative images of equatorial planes of GPMVs derived from cells expressing (A) Lck-eGFP or (B) SH4-mNeonGreen (green), together with mRuby3-tagged transmembrane domain (tm-) of CD45 (top rows, magenta) or CD86 (bottom rows, magenta; scale bar: 5 μm). Each of the latter was clustered with primary (biotin-antiHA) and secondary antibodies (streptavidin). White arrows point to sites with the signal of Lck enriched at the corresponding protein cluster.

